# Phylogenetic based dissection of eukaryotic Mo-insertase functionality: From mechanism to complex assembly

**DOI:** 10.64898/2025.12.12.693926

**Authors:** Tim Julian Schmidt, Ahmed H. Hassan, Boas Pucker, Tobias Kruse

## Abstract

Molybdenum cofactor (Moco) biosynthesis is vitally important for all organisms, yet the domain organization of the eukaryotic molybdenum insertase (Mo-insertase) remains enigmatic. We combine extensive phylogenetic reconstructions, sequence analysis and structural modeling in order to uncover evolutionary and functional principles of eukaryotic Mo-insertases. We note, that the vast majority of plant, fungi and animal species evolved fused E- and G-domains, yet the orientation of both domains in the fusion proteins differs among different eukaryotic lineages. Despite the divergent domain arrangements amongst eukaryotic Mo-insertases the E-domain active site is well conserved, with very few tolerated substitutions in >1,000 sequences. Among the Mo-insertases from different eukaryotic species, vertebrate gephyrin is the only Mo-insertase with a dual function as - next to its metabolic function - it scaffolds inhibitory neurotransmitter receptors in the post synapsis. Gephyrin is surprisingly high conserved, including surface patches not directly involved in catalysis and receptor clustering. This profile suggests additional, as yet uncharacterized, functional constraints on gephyrins evolution. Together, our results reveal how eukaryotic Mo-insertases combine evolutionary domain organization plasticity with stringent active site conservation and recognize the evolutionary constraint on gephyrińs surface conservation to be extreme, likely due to its mutual metabolic and neuronal function.

## INTRODUCTION

Molybdenum cofactor (Moco) biosynthesis is catalyzed by an ancient and highly conserved multi-step biosynthesis pathway [1], with the general steps of Moco biosynthesis being highly similar amongst eukaryotes and prokaryotes. Initially GTP is converted to cyclic pyranopterin monophosphate (cPMP), a reaction that – in all eukaryotes – was suggested to reside in the mitochondrial matrix [2]. Upon formation, cPMP is exported into the cytosol [3] where all subsequent Moco biosynthesis steps take place. In the second step of Moco biosynthesis, cPMP is converted to molybdopterin (MPT), the metal free precursor of Moco. Upon formation, MPT is used as a substrate by the molybdenum insertase (Mo-insertase). Here, notable differences exist regarding the domain organization of eukaryotic and prokaryotic Mo-insertases. Pioneering work identified *E. coli* molybdate utilization to depend on the two separate enzymes MoeA and MogA [4, 5], while the homologous domains are fused as a single polypeptide in most eukaryotes [6]. For consistency, in eukaryotes the prokaryotic nomenclature has been retained as the MoeA homologous domain of eukaryotic Mo-insertases is referred to as E-domain while the MogA homologous domain is referred to as G-domain. Work with the plant Mo-insertase Cnx1 identified the G-domain to adenylylate MPT, yielding MPT-AMP (adenylated MPT [7, 8], Fig. 1) which is used as the substrate for the subsequent molybdate insertion reaction catalyzed by the E-domain. Both Cnx1 domains form a complex which was suggested to ensure the directed and protein protected MPT-AMP transfer from G- to E-domain [9]. Upon binding to the E-domain, molybdate is incorporated into the MPT dithiolene moiety [10]. This reaction, precisely the initial molybdate binding and its transfer to the active site bound MPT (dithiolene), requires a defined set of residues which were first identified in the eukaryotic model Mo-insertase Cnx1E from the higher plant *Arabidopsis thaliana* (summarized in [6]). Consistent with the essential function of these residues for Cnx1 catalytic activity, a high degree of conservation of these residues has been reported amongst various eukaryotes [11]. Molybdate insertion into the MPT dithiolene moiety results in the formation of adenylylated Moco (Moco-AMP, [12]). Upon formation, the phosphor-anhydride bond within Moco-AMP is hydrolyzed and physiologically active Moco is released (Fig. 1, [10, 12]).

**Figure 1:**
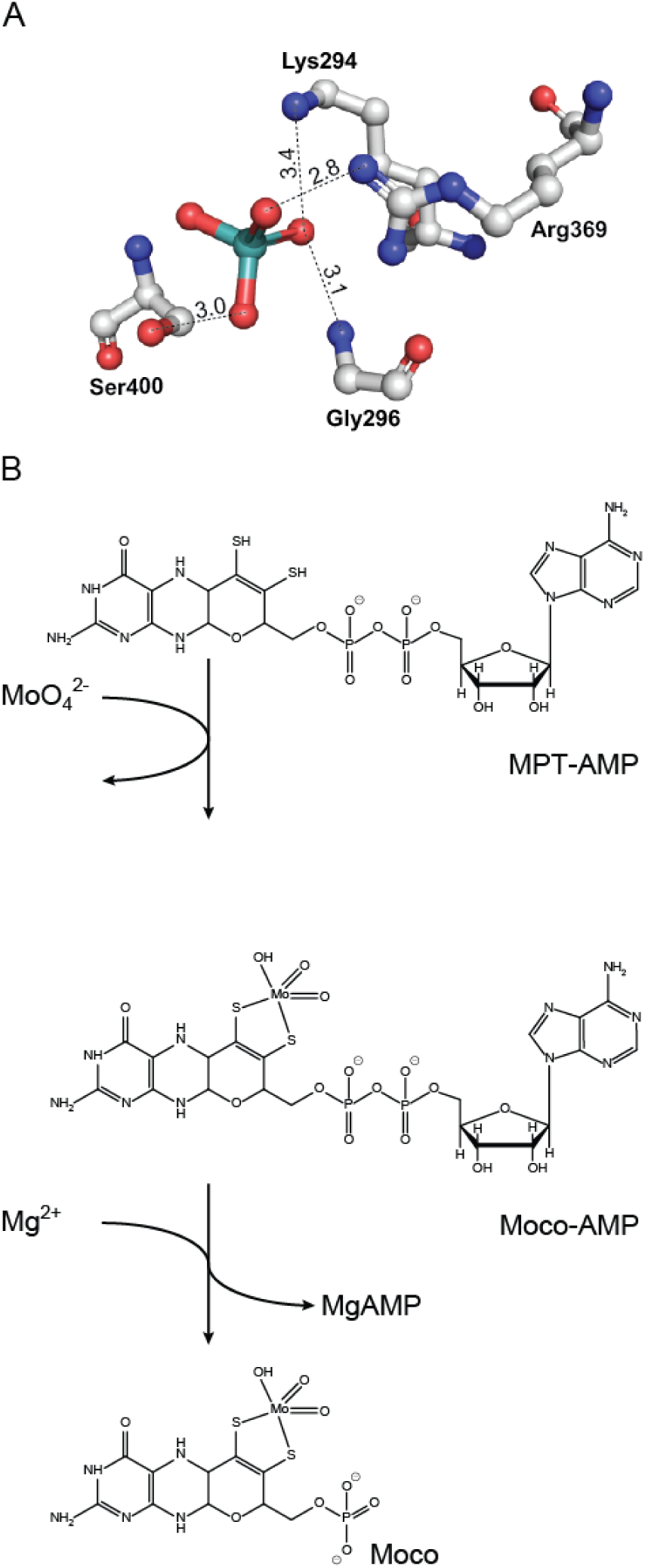
Functionalization of molybdenum. (A). Cnx1E residues involved in molybdate binding to its initial binding site [11] are shown, directed interactions are indicated by dashed lines with distances in Ångström given above; numbering refers to the *A. thaliana* Mo-insertase Cnx1E (Q39054.2 [13]). (B) The Mo-insertase G-domain catalyzes the adenylylation of molybdopterin (MPT), yielding MPT-AMP. MPT-AMP and molybdate are the E-domain substrates and most likely, a minor backbone flip within the active site results in the movement of active site bound molybdate (A) into the MPT dithiolene moiety [11] yielding Moco-AMP. A minor rearrangement of the phosphor-anhydride bond in the Moco-AMP molecule is believed to initiate its hydrolysis, resulting in the release of physiologically active Moco [10, 12].

Thus formed Moco may be transferred to Moco dependent enzymes or contribute to the celĺs insertion-competent Moco pool (summarized in [14]). Moco is the active site prosthetic group of Moco dependent enzymes (Mo-enzymes) which catalyze a diverse set of vitally important redox reactions (summarized *e.g.* in [15]). For plants, loss of nitrate reductase is most critical, as it is essentially required for plant survival [1] [16]. From the mammalian Mo-enzymes, sulfite oxidase activity is most crucial, as its depletion results in severe neurological phenotypes and ultimately leads to death of the affected individuum [17, 18]. Next to its importance for metabolic processes, Mo-metabolism is otherwise essential for mammals: Here the Mo-insertase gephyrin was first identified as a receptor clustering protein in the post synapse, but not as enzyme essential for the cellular Mo-metabolism [19, 20]. Hence explaining the name of the protein (gephyrin, greek for bridge building, [21]). Precisely, gephyrin is required for clustering of glycine- and GABA- (γ-aminobutyric acid, type A) receptors in the postsynaptic membrane of inhibitory synapses [19, 20]. Other than the plant Mo-insertase Cnx1 which forms a compact, asymmetric hexameric complex [9], gephyrin is suggested to form a lateral network which is essential for receptor clustering [22], [23] and references therein).

With very few exceptions known, eukaryotic Mo-insertases are generally assumed to possess E- and G-domains fused together in one protein, while for prokaryotes both domains exist as separate entities. To shed light on the domain organization of eukaryotic Mo-insertases in the present work we carried out an *in silico*-based approach to identify Mo-insertase sequences both in eukaryotes and prokaryotes which revealed a great number of sequences of hitherto not described putative Mo-insertases. Whilst prokaryotic Mo-insertases were identified to assemble from two separate domains, the vast majority of identified eukaryotic Mo-insertases assembled from two fused domains. Surprisingly exceptions were found in some invertebrate species, algae and protists, where both domains exist as separate entities.

All eukaryotic Mo-insertases identified within this work were found to possess a highly conserved surface patch which forms the active site. Catalytically important residues located here were found to be strictly conserved with few identified exceptions. Structural model-based analysis revealed that in six out of seven cases these will not impair functionality.

As an unexpected peculiarity, vertebrate type Mo-insertases were identified to possess a tremendously high degree of sequence conservation which - referring to the current available knowledge - is not explainable by its functions for Mo-metabolism and receptor clustering.

## RESULTS

We used the *E. coli* MoeA sequence for BLASTp searches, which allows for the identification of putative Mo-insertases from both, eukaryotes and prokaryotes. The obtained sequences were used to create a phylogenetic tree comprising a total of 332 Mo-insertases from 327 species (Fig. 2, supp. Figs. S1-S5, supplementary data files S5 and S6). For better readability, we will in the following use the term Mo-insertase instead of putative Mo-insertase throughout this work.

**Figure 2:**
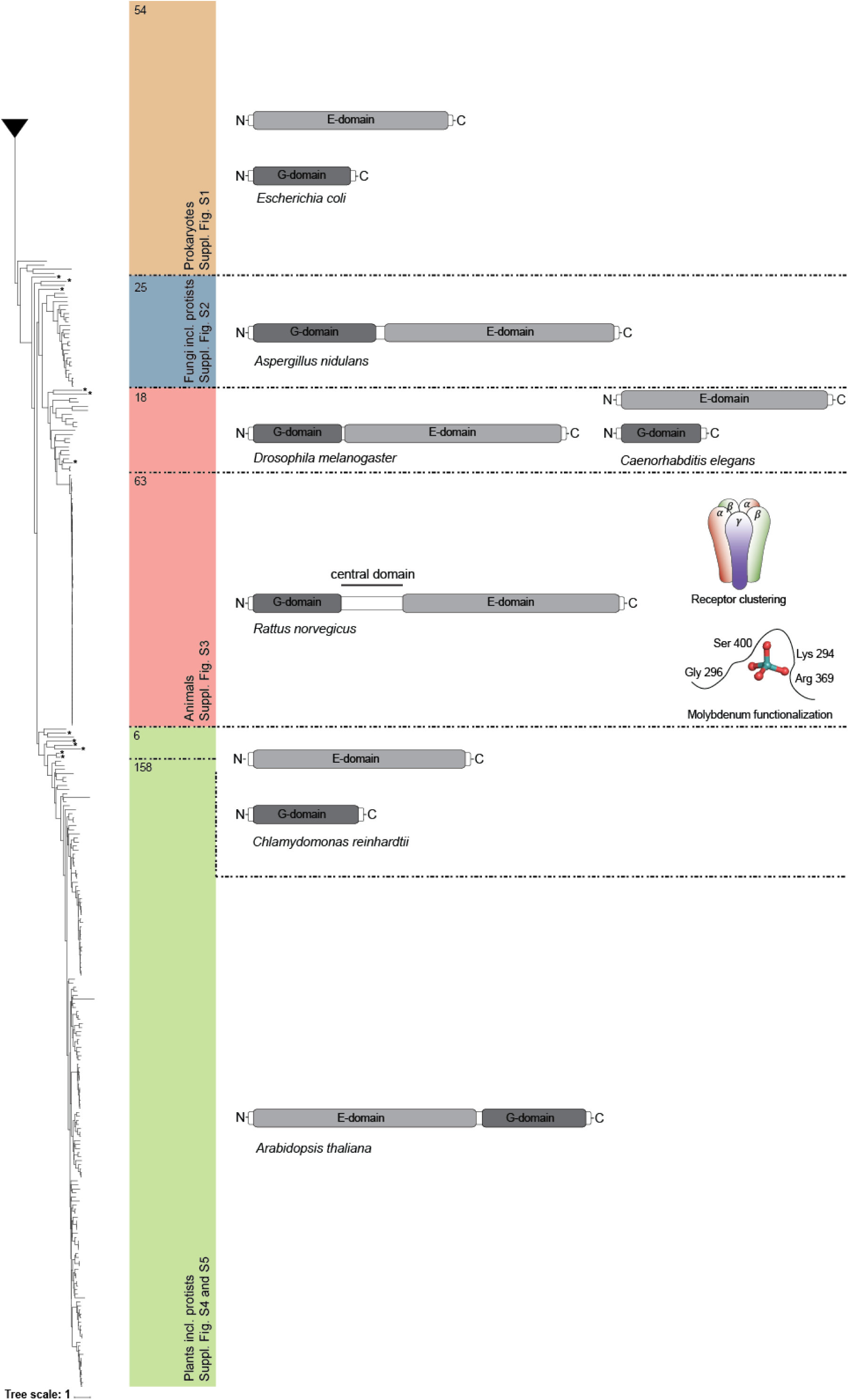
Simplified phylogenetic tree obtained from maximum likelihood analysis. The Mo-insertase families are color-coded as follows: Prokaryotic-type Mo-insertases (brownish), animal-type Mo-insertases (reddish), fungi-type Mo-insertases including protists (bluish), plant-type Mo-insertases including protists (greenish). For the detailed tree, please see supplementary figures S1 to S5 as indicated in the figure. For the sake of clarity, the prokaryotic clade has been collapsed. The dashed black lines are meant to group together Mo-insertases of different species based on their domain organization inferred within this work (see supplementary files S2, S3 and S4). As references, the domain organization of the Mo-insertases from *Escherichia coli* (MoeA: NCBI Reference Sequence NP_415348.1, MogA: NCBI Reference Sequence: NP_414550.1), *Drosophila melanogaster* (NCBI Reference Sequence: NP_726659.1, *Rattus norvegicus* (both annotations according to [13]), *Caenorhabditis elegans* (CAA90069 and CCD74267, annotation according to [24], *Aspergillus nidulans* (annotation according to [25]), *Chlamydomonas reinhardtii* (E-domain: GenBank entry DQ311646, G-domain: GenBank entry DQ311645.1, E-domain, [26]) and the model plant *Arabidopsis thaliana* (annotation according to [27]) is shown schematized. The dual function of the Mo-insertase gephyrin from vertebrates (*i.e.* clustering of γ-Aminobutyric acid type A and glycine receptors in the post synapse and functionalization of molybdenum [19, 20]) is indicated (see Fig. 1 for a detailed representation of molybdate interacting residues). The number of sequences possessing the indicated domain organization is given within the respective colored box. Asterisks indicate species which possess a differing domain organization, summarized in supplementary table S1. The tree bootstrap values are shown in supplementary file S5, the alignment file is deposited as supplementary file S6.

As expected, Mo-insertases from different taxonomic lineages group into clades, which we annotated as animal-, fungi- and plant-type Mo-insertase family respectively. On the contrary, for prokaryotes, the diversity of the identified sequences did not result in forming family-level trees (supplementary figure S1).

The animal-type Mo-insertase family comprises 84 members, which possess an N-terminal G-domain fused to a C-terminal E-domain (Fig. 2). Interestingly, exceptions were identified for the Mo-insertases from invertebrates namely *Trichuris trichiura* (*T. trichiura)*, *Caenorhabditis elegans* (*C. elegans*) and *Pinctada imbricata* (*P. imbricata*), supplementary table S1. In these organisms, E- and G-domain were found to occur separately.

The fungi-type Mo-insertase family comprises in total 28 members including unicellular organisms from the SAR clade (*i.e. Tribonema minus*, *Symbiodinium necroappetens* and *Reticulomyxa filosa*) and the zooflagellate *Tecamonas trahens*. We found the vast majority of fungal Mo-insertases identified in this work to possess an N-terminal G-domain and a C-terminal E-domain (Fig. 2 and supplementary figure S2). However, few exceptions were identified, summarized in supplementary table S1, figure S1 and figure S6).

The vast majority of plant Mo-insertases is assigned to members of the *Streptophyta* phylum and possess an N-terminal E-domain fused to a C-terminal G-domain (see Fig. 2 for comparison). A few *Streptophyta* were identified to harbor a separate E-domain (supplementary table S1) however, here no G-domain was identified which may be best explained by a provisional and/or incomplete annotation in the employed databases. Several unicellular organisms are summarized in supplementary table S1 are part of the plant type Mo-insertase group, including the alga *C. reinhardtii* for which was reported, that here the E- and G-domains are expressed separately ([26], see Fig. 2 for comparison). We identified this to hold true for all unicellular organisms included into the plant-type Mo-insertase group (supplementary table S1).

Having identified previously undescribed putative Mo-insertases in animals, fungi and plants, we next went on to confirm that these are indeed *bona fide* Mo-insertases. Therefore positional homologs to *A. thaliana* critical active site residues [6] were identified as described previously (detailed in the material and method section and [28]; table 2, table 3).

**Table 2:** Conserved active site residues of the eukaryotic Mo-insertase E-domain. The catalytically important residues of *A. thaliana* Cnx1E (Cnx1 sequence NP_197599.1) have been tabulated. Positional homologs of invertebrates, vertebrates, fungi and plants have been identified and the number of identical residues as compared to the Cnx1 active site residues has been calculated as rounded percentage. Non-conserved residues are likewise included and the rounded frequency (percentage) with which these occur is given. In total, 21 invertebrate-type, 62 vertebrate-type, 26 fungi-type, and 155 plant-type Mo-insertases were considered. X = alignment gap.

| Annotation | *A. thaliana* residue |  |  |  |  |  |  |
| --- | --- | --- | --- | --- | --- | --- | --- |
|  | Frequency [rounded %] |  |  |  |  |  |  |
| *Arabidopsis thaliana* | D274 | K294 | G296 | K297 | S328 | R369 | S400 |
| Invertebrates | 100 | 100 | 100 | 5<br>L 95 | 100 | 100 | 100 |
| Vertebrate | 100 | 100 | 100 | 0<br>L 100 | 100 | 100 | 100 |
| Fungi | 100 | 96<br>R 4 | 96<br>A 4 | 96<br>R 4 | 96<br>T 4 | 96<br>K 4 | 96<br>T 4 |
| Plants | 99<br>X 1 | 92<br>R 8 | 100 | 100 | 99<br>X 1 | 99<br>X 1 | 99<br>X 1 |

**Table 3:**
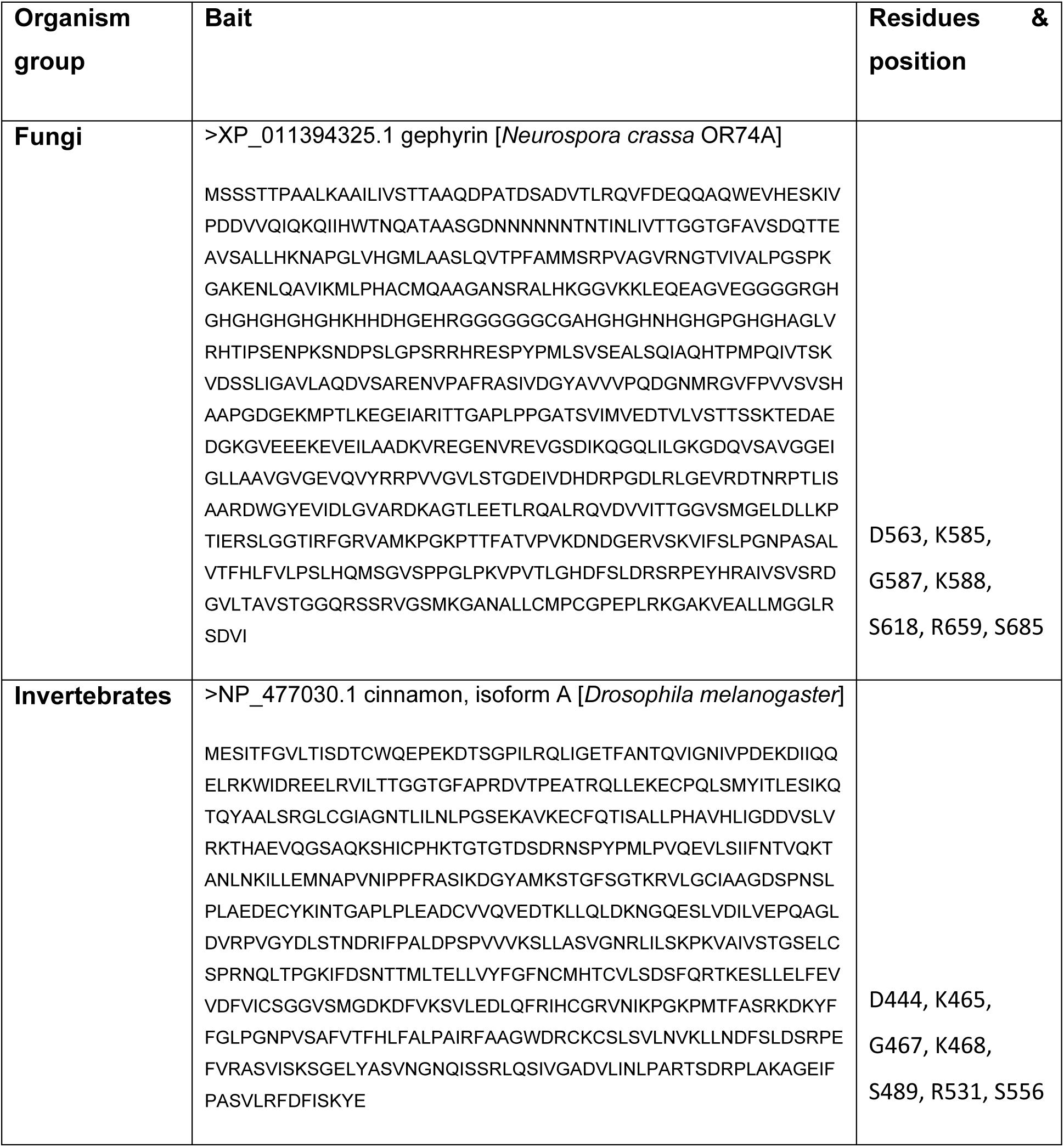
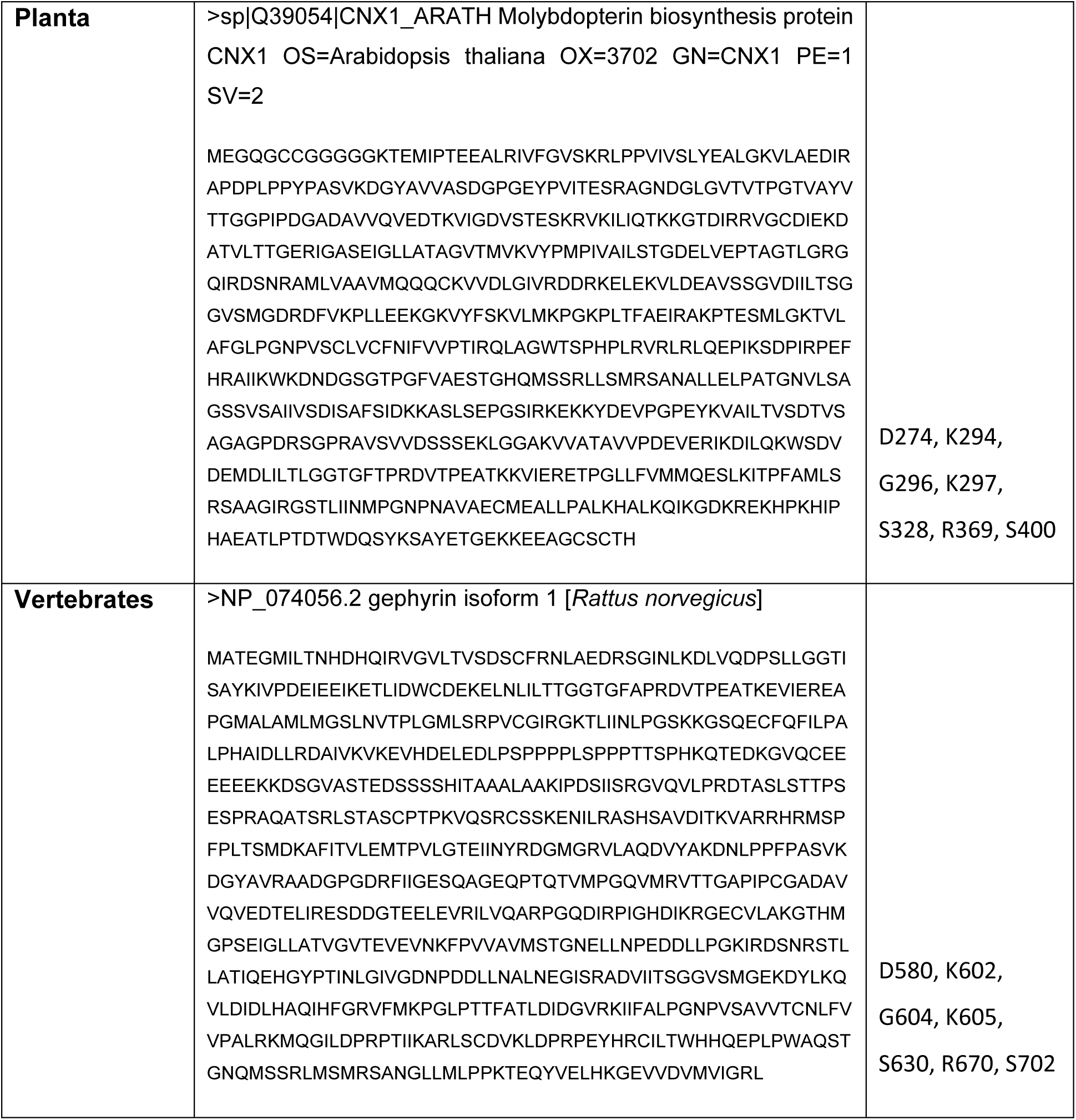
Reference sequences, residues and their positions for the inspection of active site amino acid residues in Mo-insertase candidates. .

To elucidate the potential impact of the identified non conserved residues on Mo-insertase functionality we carried out an *in silico*-based approach in which the structure of respective Cnx1E variants was predicted using AlphaFold [29, 30] (Figure 3). Modelling of all variants was possible (see also suppl. Fig. S9 for comparison), however for variant G296A, only the “relaxed” backbone conformation present in 6ETF [11] yielded a viable template for modeling.

**Figure 3:**
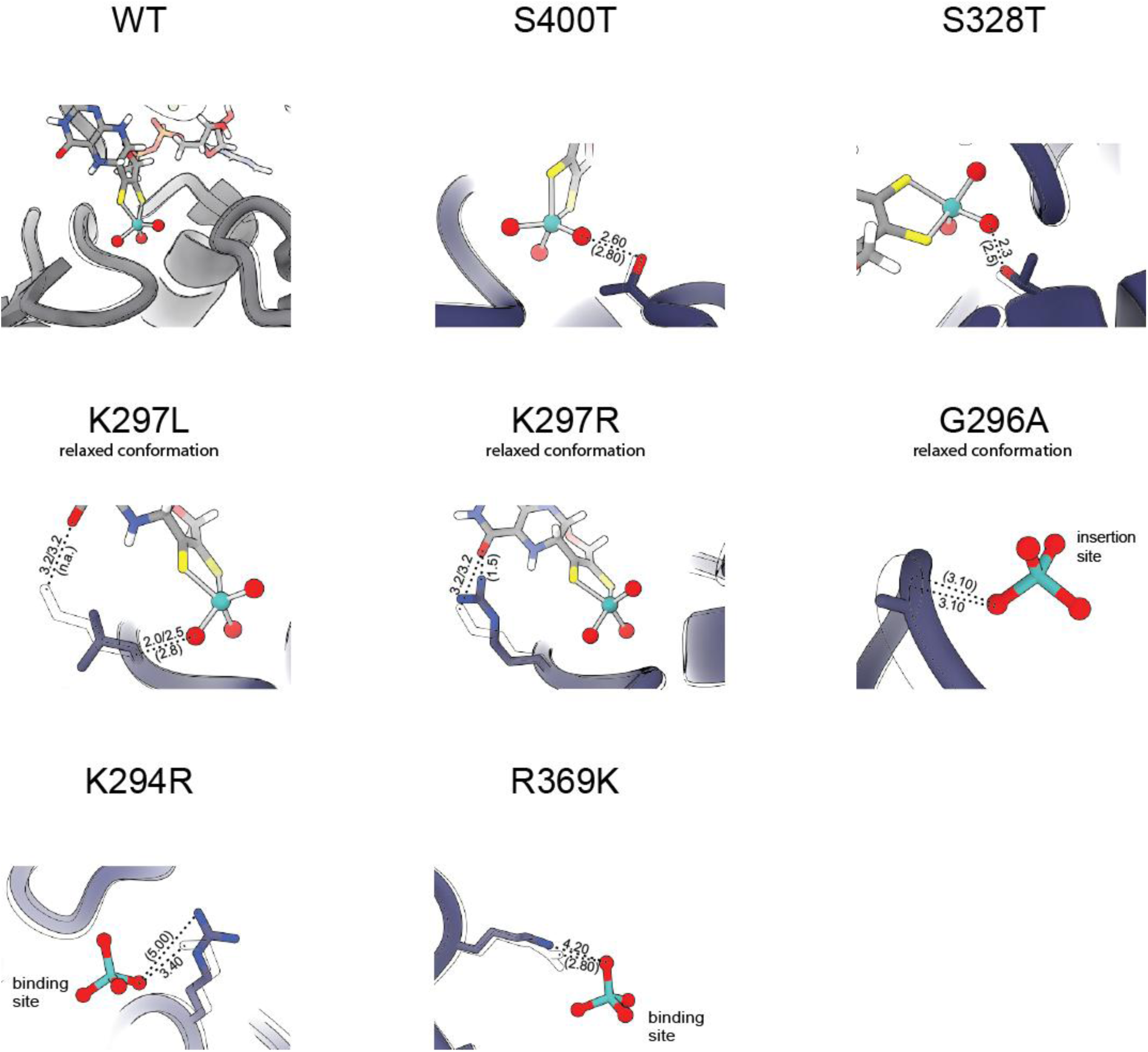
AlphaFold-Based modelling of Mo-insertase active site variants. Cnx1 variants possessing the active site variations identified and tabulated in table 2 were modelled using AlphaFold (as specified in the material and methods section). View of the active site structures of the modelled Cnx1 variants. Cnx1 variants K297R/L, S328T, S400T: Moco-AMP is shown derived from the structural superimposition of modelled variants with protein structure 6Q32 [12]; Cnx1 variants G296A, K294R and R369K: Molybdate is shown derived from the structural superimposition of modelled variants with protein structure 6ETF [11]. The modelled active site structures are shown in ribbon representation; exchanged residues are depicted as sticks and colored blue. The outlines of the respective positional homologous wildtype residues were derived from the structural superimposition of modelled variants with the protein structure 6ETF [11] and are shown superimposed. Moco-AMP and molybdate are shown in ball and stick representation. Dashed lines indicate directed interactions between modelled and non-modelled active site residues with Moco-AMP and/or molybdate respectively, with distances given in Ångström (Å). Brackets indicate distances between modelled residues, while distance measurements involving wildtype residues are given without brackets.

Within the animal-type Mo-insertase family, a large number of sequences groups together extremely close (detailed in Fig. 4). Interestingly, these are all assigned to jawed vertebrates (*Gnathostomata*) where the Mo-insertase (named gephyrin here, [21]) is known to possess a neuronal function next to its function in Mo-metabolism.

**Figure 4:**
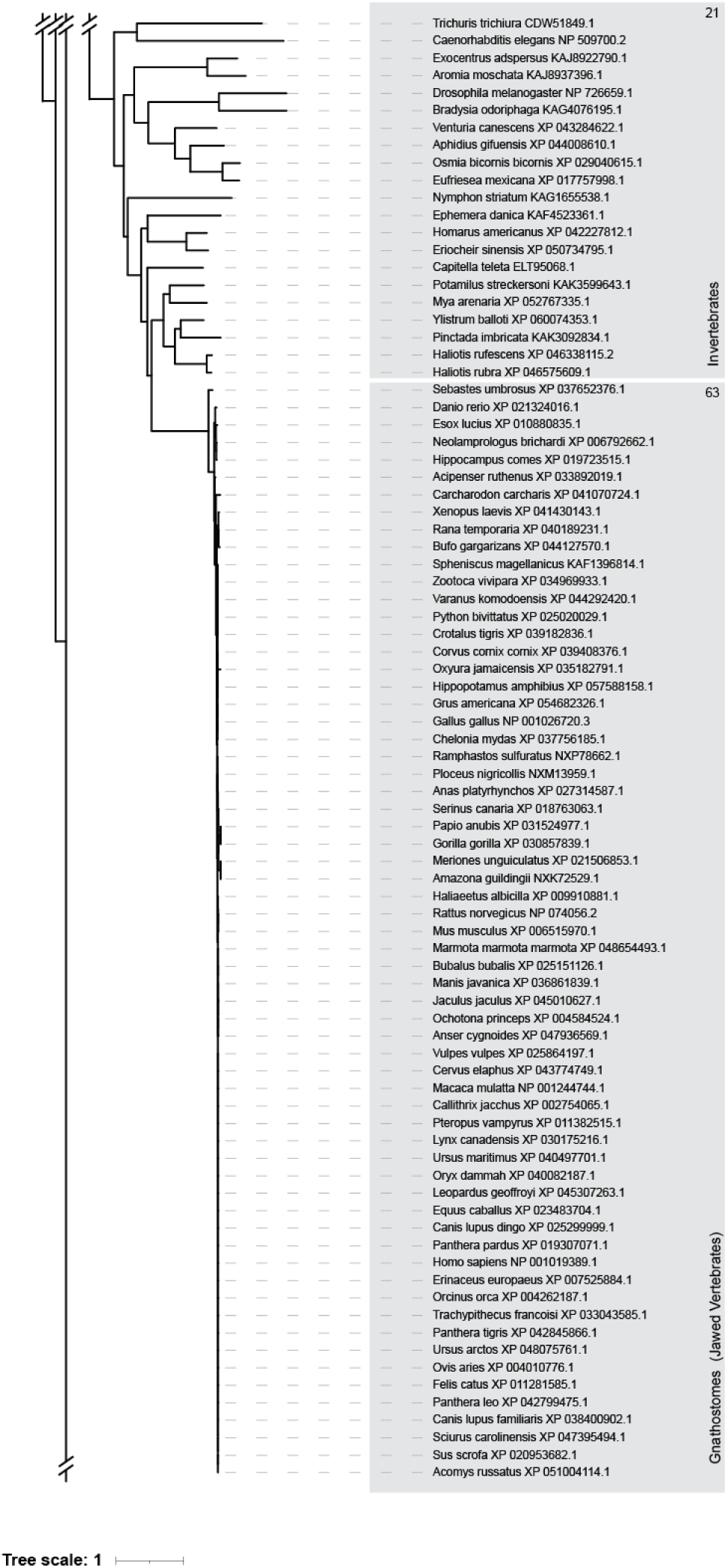
Phylogenetic distance tree of animal-type Mo-insertases. Supplementary figure S2 has been modified and is shown here. Mo-insertases identified in invertebrates possess sequence deviations which result in their arrangement within the phylogenetic tree allowing it to assign a phylogenetic affiliation *i.e. Nematodes*, *Insecta* and *Mollusca*. Jawed vertebrates Mo-insertases do not arrange group-wise, hence preventing the assignment of any phylogenetic affiliation. Clades harboring plant and fungal orthologs have been omitted in this figure due to space constrains (see supplements for details). Numbers refer to the number of sequences forming the respective groups.

We are fully aware, that this extremely close grouping – at first sight – may suggest corrupted database entries or subsequent processing errors. While this might appear as an artifact, all our investigations support the validity of these sequence records.

To exclude any influence of the phylogenetic age of the vertebrate trait on the degree of gephyrińs sequence conservation, we next analyzed the patristic distances between species in the taxon *Gnathostomata* (jawed vertebrates, Fig. 5) and compared these with values observed for taxa of similar age, documenting almost no changes in gephyrin during *Gnathostomata* evolution. To further substantiate this finding we next went on and constructed two additional protein phylogenies as controls, one for the Moco biosynthesis enzyme MOCS2B [31], (supplementary file S7) and one for alcohol dehydrogenase ([32], (supplementary file S8) and analyzed the respective patristic distances (supplementary figure S6). As can be seen from supplementary figure S7, the patristic distances of ADH sequences in the taxon *Gnathostomata* (1.67) are higher than those of gephyrin (0.04). The MOCS2B protein phylogeny revealed a recent splitting as two large metazoan protein families *i.e. Amphibia/Dinosauria* and *Mammalia* were identified, while no *Gnathostomata* trait as observed for gephyrin / ADH were observed. Comparison of the patristic distances of both families again revealed a higher patristic distance, *i.e.* 0.94 for *Amphibia/Dinosauria* and 0.45 for *Mammalia*, respectively. Hence even within this – as compared to *Gnathostomata* (462 MYA, Fig. 5) - younger taxa (*i.e.* ca. 300 MYA, *Amphibia/Dinosauria* and ca. 180 MYA, *Mammalia*) a significantly higher patristic distance was identified for an enzyme from the same biosynthesis pathway.

**Figure 5:**
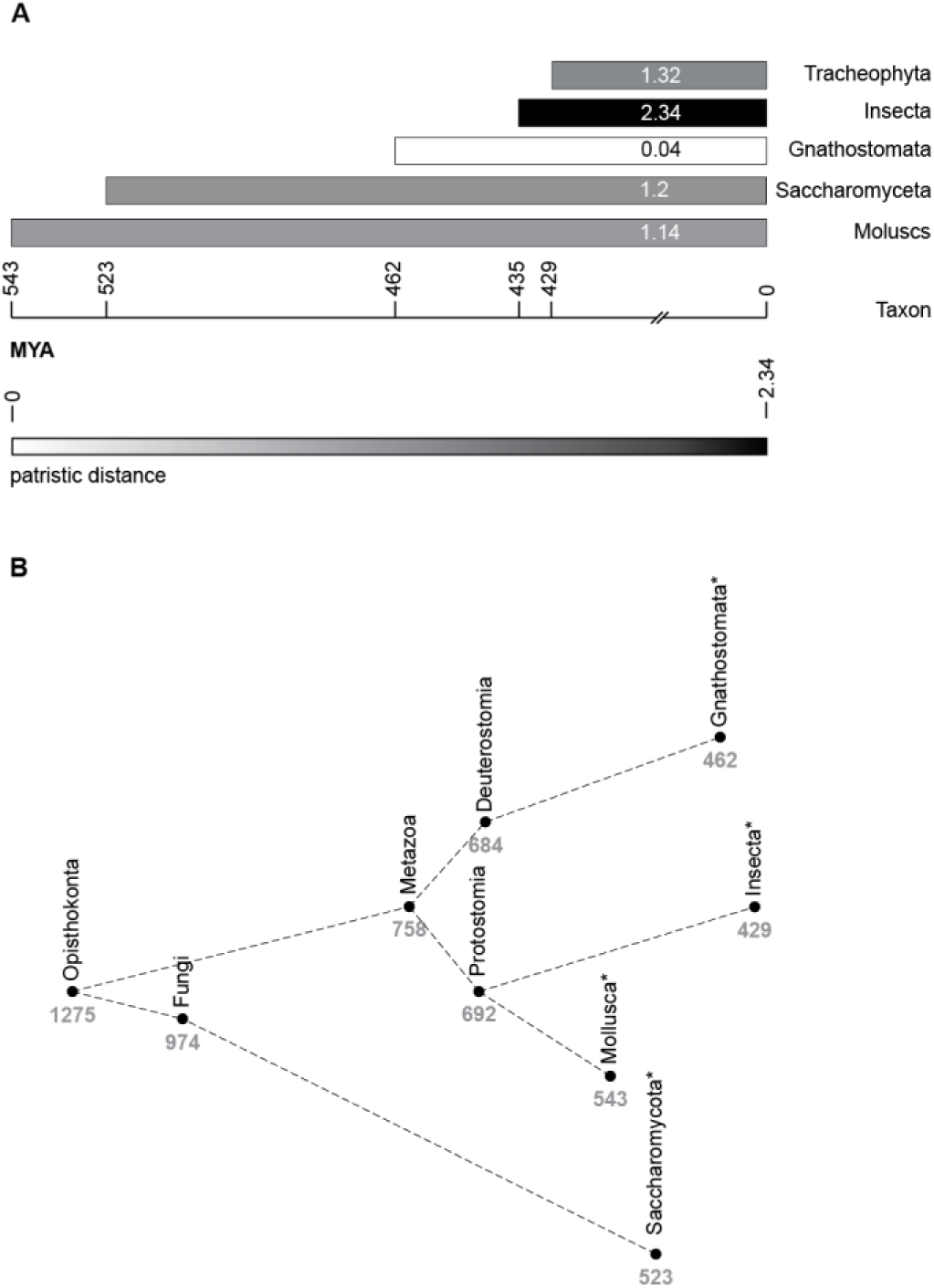
**Average patristic distance between Mo-insertases in different taxa. (**A) Patristic distance of the Mo-insertases within indicated taxa. The estimated age of the compared taxa is indicated (mya = million years ago). Colors correlate with the patristic distance as indicated within the figure. (B) Simplified timetree visualizing the phylogenetic relationship of selected eukaryotic non-plant taxa. The numbers specify the estimated taxon-ages. Asterisks indicate the taxa compared in (A). (A) and (B): Estimates of taxon ages were extracted from the evolutionary time tree of life [33, 34].

In summary, our patristic distance analysis documents that the highest degree of gephyrins sequence conservation is not a result of the phylogenetic age of the vertebrate trait.

In the following known residues that are important for the two functions of gephyrin (*i.e.* receptor clustering and Moco synthesis) were identified and plotted onto the protein surface. As can be seen from Fig. 6, a significant part but not the complete protein surface is associated with these functions while the overall surface conservation was found to be very high (supplementary Figure S8).

**Figure 6:**
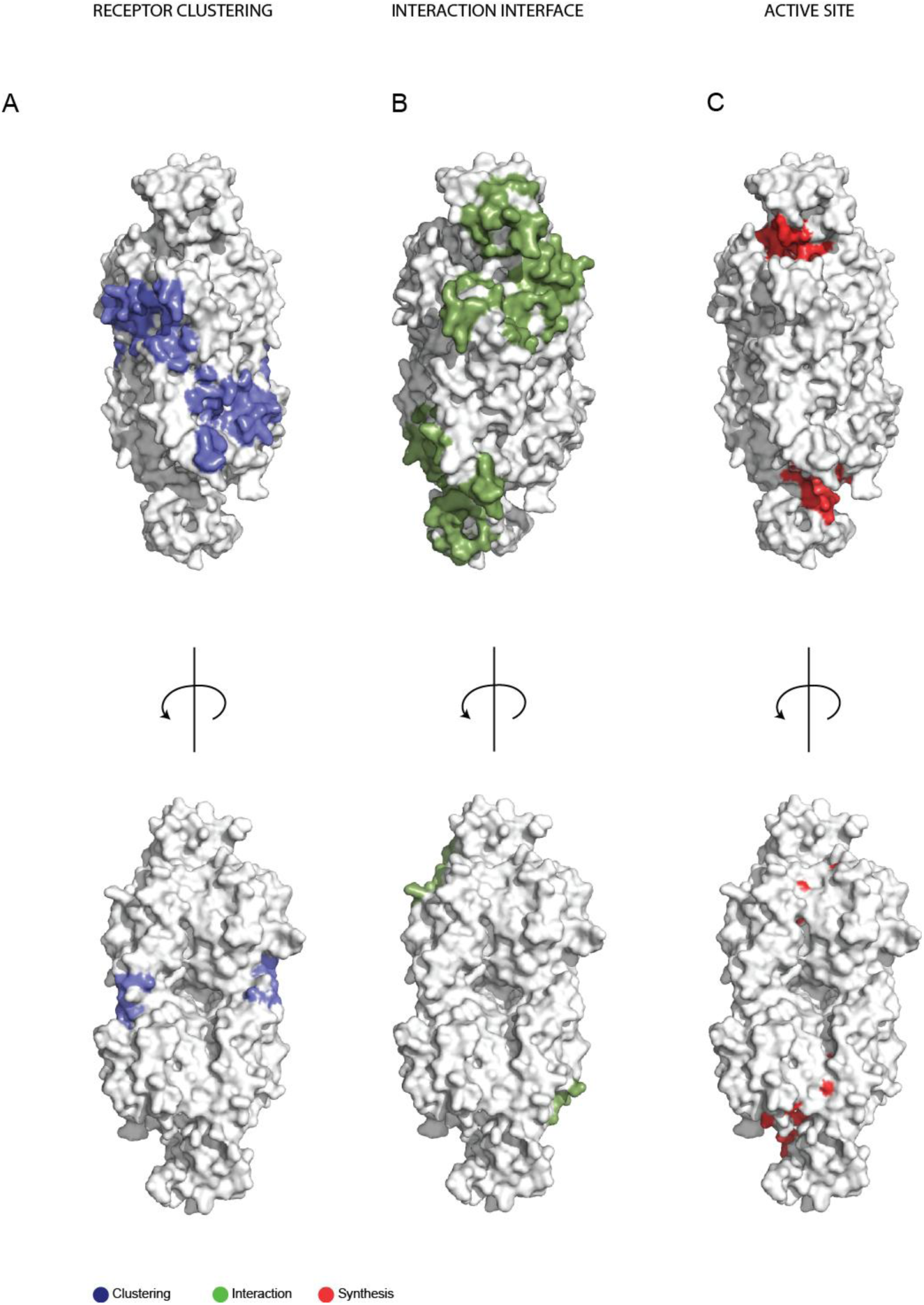
Functionally relevant residues of the gephyrin E-domain. Surface representation of the mammalian Mo-insertase E-domain (PDB code: 2FU3). (A) Residues involved in receptor binding (according to [35]) are shown in blue. (B) Interfacing residues of the interaction 2 model of the plant Cnx1 Mo-insertase complex [9] are shown in green. (C) Residues involved in molybdenum functionalization of the plant Mo-insertase Cnx1 are shown in red (recently summarized in [6]).

## DISCUSSION

Within this work we carried out an *in silico*-based approach to identify Mo-insertases in the species of life. Our work revealed in total 332 Mo-insertases from 327 (in total) eukaryotic and prokaryotic species. The finding that for prokaryotes the diversity of the identified sequences did not result in forming family-level clades may be best explained by the prevalent horizontal gene transfer here [36] leading to conflicting gene histories [37]. However, as known for the model Mo-insertase MoeA from *E. coli*, the prokaryotic Mo-insertases identified all assemble from a separate E- and G-domain.

Relating to the domain organization, our work classified all novel eukaryotic Mo-insertases identified either as plant-type (N-terminal E-domain, C-terminal G-domain) or animal/fungi-type Mo-insertases (N-terminal G-domain, C-terminal E-domain), with the notable exceptions of Mo-insertases identified in some protists, algae and invertebrates which – like prokaryotes – all possess the E- and G-domain as separate entities (as documented by 190 manually carried out and inspected pairwise sequence alignments (see supplementary data files S2, S3 and S4) and when indicated *in silico* based complementary structure predictions, see supplementary table S1).

Very recently, Megrian and colleagues [38] described the fusion of E- and G-domain to be present in all studied animal type Mo-insertases which they suggested to be due to the finding that this is mandatory for gephyrińs neuronal function (*i.e.* receptor clustering in the post synapse of inhibitory neurons). Our study revealed that in some animals (*i.e. C. elegans* (see also [24]), *T. trichuria* and the mollusk *P. imbricata,* see supplementary table S1 for comparison) the Mo-insertase E- and G-domains exist as separate entities, hence the tendency of E- and G-domain to occur fused together in animals requires a careful review. As the invertebrate-type Mo-insertases reported within this work essentially lack conserved E-domain residues required for receptor interaction (supplementary Figure S10), we reason, that any vertebrate-like neuronal function of invertebrate-type Mo-insertases is unlikely, which is consistent with the lack of critical residues for receptor clustering in *C. elegans* and *Drosophila* Mo-insertases [39].

Why do eukaryotes possess inconsistent E-G domain arrangements? Other than eukaryotes, all prokaryotes studied thus far possess the identical domain arrangement: E- and G-domains are separate entities, documenting that domain fusion is not mandatory for Mo-insertase functionality (here). Interestingly, when recombinant plant Cnx1E- and G-domains are expressed separately, the MPT-AMP transfer from G- to E-domain as well as the subsequent metal insertion reaction occur under fully defined *in vitro* conditions when both enzymes are co-incubated [10, 12, 27]. Further, the *Chlamydomonas reinhardtii* E-domain was found to complement an *E. coli moeA* mutant strain [40]. These findings point towards, that i) eukaryotic Mo-insertase functionality does not mandatorily depend on domain fusion and ii) metabolite transfer from G- to E-domain likely underlies a common principle in eukaryotes and prokaryotes (see also [40]).

The reason for domain fusion in eukaryotic Mo-insertases may be best explained by assuming that this allows for a directed [9] and efficient MPT-AMP transfer from G- to E-domain [41], which may be beneficial for the vast majority of eukaryotic species reported within this work (see also [38] for comparison).

It is not clear, when the E- G- domain fusion occurred, however it appears to have occurred independently at least twice during the evolution of eukaryotes, documented by the finding that plants possess an inverted domain orientation as compared to animals and fungi [25, 42]. We suggest that the identified separate E- and G-domains in *C. elegans*, *T. trichuria* and the mollusk *P. imbricata* (see also supplementary table S1) may result from the division of the fused domains during speciation. The other possible explanation – an independently occurred domain fusion during speciation of the various animal species – appears unlikely. However, the findings about domain fusion prevalence and evolutionary conservation are based on the sequence data that is currently available and may be improved as more genome sequences and derived polypeptide sequences become available.

According to our analysis, a number of catalytically significant residues for Mo-insertases of the plant, fungal, and invertebrate types are conserved across species, which is in line with their suggested functional roles. For the exceptions identified we carried out AlphaFold based modelling to deduce the impact the identified variation may have on E-domain functionality. Based on our modelling approach (see Fig. 3), we reason that the identified alterations S400T and S328T most likely will not impact functionality, as the Ser OH-group is essentially “replaced” by the threonine OH-group within the active site and no obvious impact of threoninés methyl group on active site chemistry could be identified. The exchange of Lys 297 to Arg or Lys 297 to Leu will not have any impact on the interaction with the Mo-center: It is the amide nitrogen atom which is involved in a directed interaction here [11]. However in Cnx1E variant S269D D274S (co-crystallized with Moco-AMP, [12]), the side chain of K297 is involved in a single, directed interaction with the Moco-AMP pterin part. For variant K297R our modelling revealed that a directed interaction likely will occur, while our modelling data suggests, that this will not be the case for variant K297L. However, obviously this does not impair functionality, as all vertebrate type Mo-insertases and 95% of the invertebrate type Mo-insertases reported here (Table 2) possess the K297L variation. Cnx1E residues K294 and R369 are involved in (initial) molybdate binding [11] [12] to the active site. Exchange of Lys 294 to Arg or Arg 369 to Lys will very likely have no impact on the (positive) charge of the binding site which leads us to the assumption that binding of molybdate is most likely not impaired in the respective variants. Our modelling suggests that solely the G296A variant likely will possess an impaired functionality: The Cnx1E G296-K297 segment exists in two conformations [11] from which variant G296 cannot adopt the “tensed” conformation as documented by the finding that modelling with the respective template was not possible. As (initial) molybdate binding is linked to the “tensed” conformation [11], we conclude that G296A is impaired in molybdate binding which most likely will exclude the subsequent molybdate insertion. From all Mo-insertases identified and characterized thus far, vertebrate type Mo-insertases possess an extraordinarily high degree of sequence conservation (see also supplementary Fig. S8B). As other vertebrate enzymes such as ADH and MOCS2B possess a significantly lower degree of sequence conservation (documented by the respective patristic distances, see Fig. 5), we conclude that the evolutionary rate of the bifunctional gephyrin is lower. We note, that gephyrin residues involved in its metabolic (Mo-functionalization and complex formation) and receptor-clustering function locate nearly exclusively to the “front-sidé side of the enzyme, while merely no known functional relevant residue(s) locate to the “back-sidé (Fig. 7).

**Figure 7:**
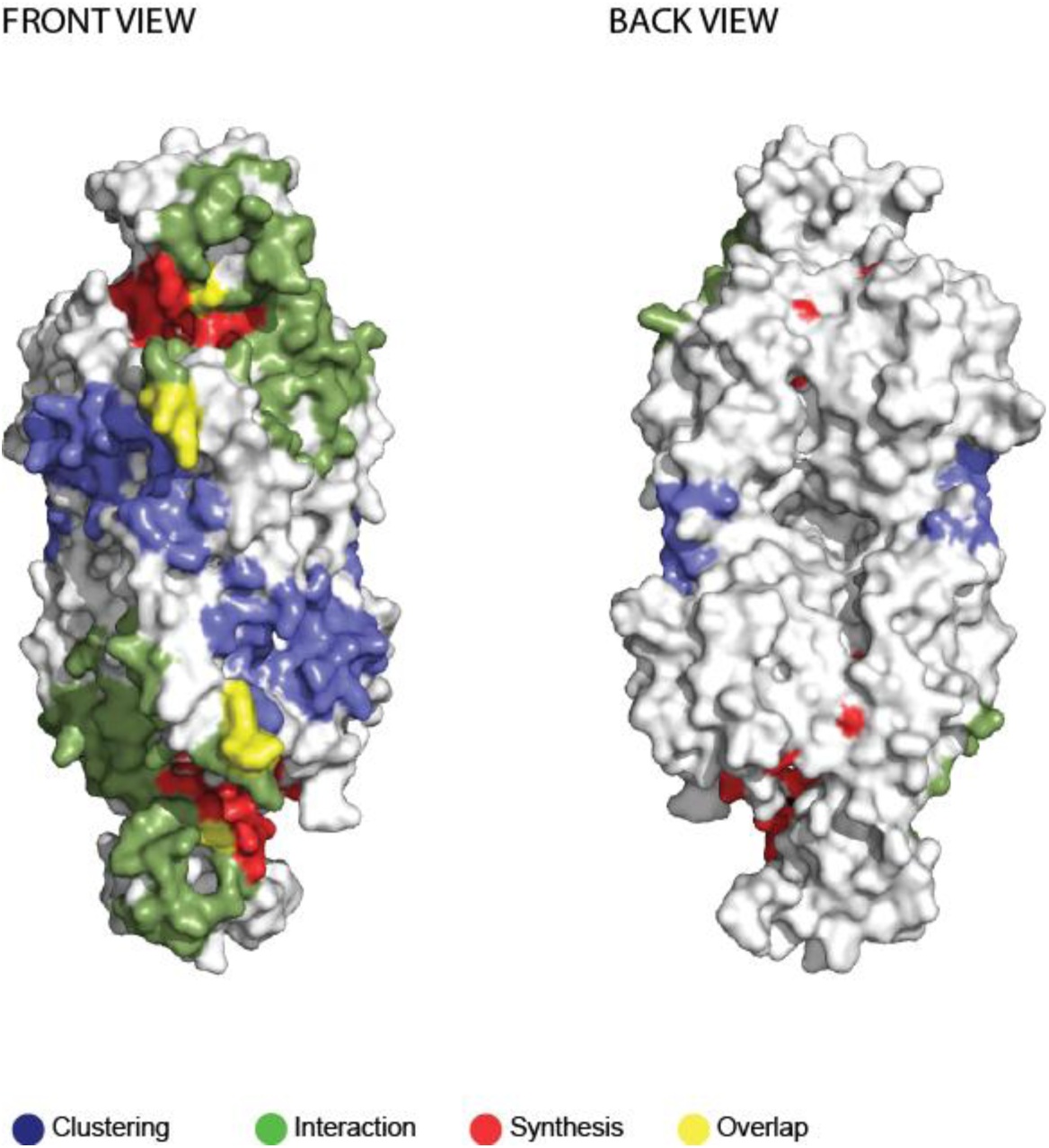
Functional relevant residues of the gephyrin E-domain. All functional relevant residues (clustering, interaction, synthesis) shown individually in Fig. 6 are shown in a combined image here. Please see legend of Fig. 6 for details. Residues shown in yellow result from an overlap of the functional relevant residues.

We speculate that putatively the remaining conserved surface area (see also supplementary Figure S8B) is crucial for other known evolutionary conserved protein interactions or post-translational modifications see *e.g.* [45]. Given the fact that gephyrin forms high molecular weight complexes beneath the post synaptic membrane a function to gephyrińs backside may relate to this.

## MATERIAL AND METHODS

*Retrieval of Bait and Reference Sequences –* In an initial setup, the protein sequence of *Escherichia coli* MoeA (WP_003903624.1) was used as query in a BLASTp search see Figure 8 for an overview.

**Figure 8:**
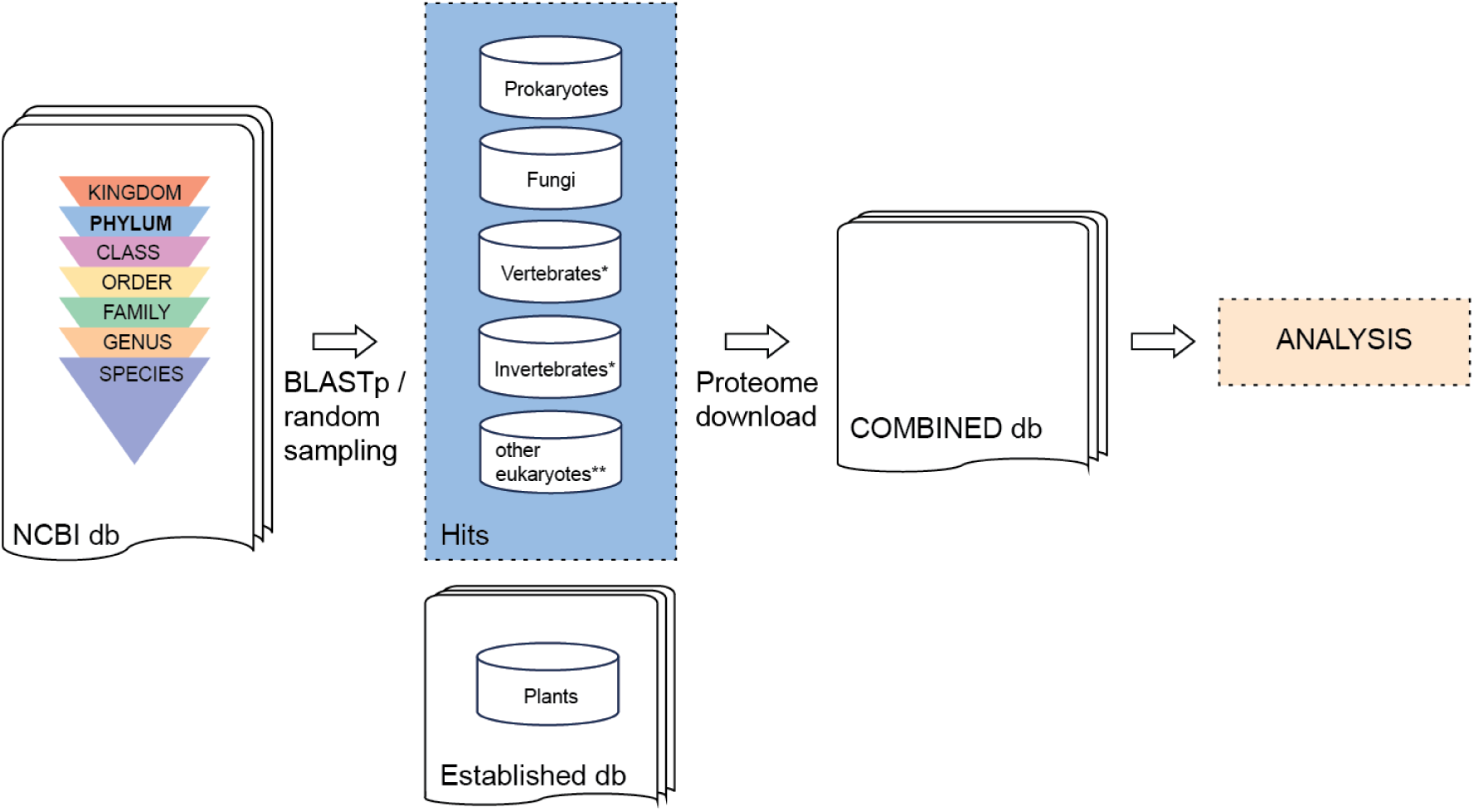
Simplified taxon sampling scheme: Essentially, the MoeA protein sequence was used as query in BLASTp searches against the NCBI database (db). Doing so revealed numerous prokaryotic and eukaryotic phylae to contain MoeA homologs from which the corresponding proteomes were downloaded and included into a new, combined db. *For vertebrate and invertebrates a random sampling approach was chosen which allowed it to include at least one representative proteome per phylum. **The taxonomic groups included into the combined db from “other eukaryoteś are described in detail in the following section. A readily established plant db was likewise included into the combined db. If not stated otherwise, subsequent analysis was carried out exclusively with this established combined db. Please see the following sections for details.

Initially, BLASTp v2.15 searches were carried out, using the NCBI protein database (non-redundant protein sequences (nr) with default settings between March and June 2024. Initially, the top two hits (referring to the bit score of the NCBI BLASTp search result) for all bacterial phylae were identified. Next the annotated polypeptide sequences of the respective species (see Data file S1) were downloaded from the NCBI database [46]. In few cases, no annotated polypeptide sequences were available for the species of the top two hits obtained from BLASTp searches. Here, the first two hits for which annotated polypeptide sequences were deposited were considered. Thus obtained sequences were used for downstream analysis. For fungi, MoeA (WP_003903624.1) was used as query in BLASTp searches using the NCBI protein database (non-redundant sequences (nr), default settings) in the following taxonomic groups: *Ascomycota, Basidiomycota, Dikarya incertae sedis, Blastocladiomycota, Chytridiomycota, Cryptomycota, Microsporidia, Mucoromycota, Nephridiophagidae, Olpidiomycota, Sanchytriomycota,* and *Zoopagomycota*. Referring to the NCBI taxonomy browser [47] *Ascomycota, Basidomycota, Chytridiomycota, Cryptomycota, Microsporidia, Mucoromycota, Olpidiomycota, Sanchytriomycota* and *Zoopagomycota* are fungal phyla, while *Nephridiophagidae* is a fungal subclass ranked as phylum and *Dikarya incerta sedis* are unranked species. From each taxonomic group, again the top two hits (referring to the bit score of the BLASTp search result) were identified and the annotated polypeptide sequences were downloaded from the NCBI database [46] and used for downstream analysis. As we sought to use plants as a control group, we decided to use an available dataset comprising the annotated polypeptide sequences of in total 147 plant species (supplementary data file S1). For animals, the annotated protein sequences of well characterized model organisms *i.e. Mus musculus*, *Gallus*, *Rattus norvegicus*, *Danio rerio*, *Xenopus laevis*, *Drosophila melanogaster*, *Caenorhabditis elegans* and that of *Homo sapiens* were downloaded from the NCBI database [46] and used for downstream analysis. Upon identification of hits with highest sequence similarities amongst the analyzed jawed vertebrate (*Gnathostomata*) species, the number of jawed vertebrate species included into analysis was significantly expanded to cover various species from different clades aiming to obtain a comprehensive dataset. Therefore, an extension of the animal dataset was based on a BLASTp v2.16 analysis against nr with the same settings in January 2025. In total the analyzed vertebrate dataset encompassed 63 sequences from 59 geni and 63 species, respectively. The annotated polypeptide sequences of selected *Gnathostomata* species were downloaded from the NCBI database [46] and used for downstream analysis. As highest Mo-insertase sequence similarity was only identified amongst (jawed) vertebrate species, we sought to analyze Mo-insertase sequences from invertebrate species for comparison. As compared to initial taxon sampling, the number of invertebrate species included into analysis was significantly expanded to cover various species from the phyla *Nematoda* and *Mollusca* and the class *Insecta* [47]. Annotated protein sequences of selected invertebrate species were downloaded from the NCBI database [46] and used for downstream analysis. Finally, the MoeA sequence (WP_003903624.1) was used as query in BLASTp searches targeting eukaryotic taxons not covered by the hitherto described approaches, thus identifying MoeA related sequences in various protists and algae. From each taxonomic group [47], precisely *Amoebozoa* (clade), *Ancyromonadida* (clade), *Apusozoa* (class), *Breviatea* (class), *Dephylleia* (genus), *Regifilida* (order), *Mantamonadidae* (genus), *Cryptophyceae* (class), *Discoba* (clade), *Glaucocystopheae* (class), *Haptophyta* (phylum), *Centroplasthelida* (class), *Malawimonadida* (order), *Metamonada* (clade), *Opisthokonta* (class), *Rhodophyta* (phylum), *Alveolata* (clade), *Rhizaria* (clade), *Stramenopiles* (clade), *Telonemia* (genus), *Kathablepharidaceae* (order), *Palpitomonas* (genus), *Virdiplantae* (kingdom), *Ancoracysta* (genus), *Picozoa* (genus) and *Hemimastigophora* (phylum), the top two hits (referring to the bit score of the NCBI BLASTp search result) were identified and the respective, annotated polypeptide sequences were downloaded from the NCBI database [46] and used for downstream analysis.

*Identification of gephyrin homologs in Cyclostomata* – For identification of gephyrin homologs in *Cyclostomata*, a BLASTp analysis with the *H. sapiens* gephyrin sequence (Q9NQX3.1 [48]) as query against the NCBI protein sequence database (restricted to the clade *Cyclostomata* and using non-redundant protein sequences (nr), default settings) [49] was carried out. Again, the top two hits referring to the bit score of the NCBI BLASTp search were identified and the identified protein sequences (*i.e.* XP_032821144.1 and XP_061424737.1) were used to prepare an alignment using Clustal Omega [50].

*BLAST* – The *Escherichia coli* MoeA sequence (WP_003903624.1) was blasted against all downloaded polypeptide sequences using the script collect_best_BLAST_hits.py and default settings [51]. The obtained sequences were subsequently used for phylogenetic analysis.

*Phylogenetic analysis* – Generation of alignments was carried out using MAFFT v7.526 [52]. Phylogenetic trees were constructed with IQ-TREE2 v2.3.4 [53] using the maximum likelihood estimation and 1000 bootstrap replicates. The model used was LG+R8 (MoeA and ADH-trees). For the MOCS2B tree, the model used was JTTDCMut+R7. ITOL v6 was used to visualize the phylogenetic trees [54]. Patristic distances within clades were calculated using the Python script branch_length_comparison.py (https://github.com/bpucker/molyb) based on the dendropy module [55].

*Elimination of contamination and splice variants in the dataset* – After initial tree building, protein sequences from the same species, possessing highest sequence similarities, have been manually inspected for obvious annotation errors (eukaryotic and prokaryotic sequences) and splice variants (eukaryotic sequences). Given that protein sequences originating from multiple splice forms were identified, subsequent analysis was carried out with a single sequence per species. Here the top hit obtained from the BLAST using the specified script (see above) was used for further analysis.

Afterwards, the dataset was inspected manually a second time, to identify species harboring more than one protein sequence as identified by BLASTp analysis. Identified sequences were then checked for their origin to rule out contaminations using BLASTp. Thus identified contaminants were tabulated in supplementary table S2.

When more than one sequence per species remained in the data set upon application of the above-described regime, routinely all sequences from a single species were aligned to the *Escherichia coli* MoeA protein sequence (WP_003903624.1) and the *Homo sapiens* gephyrin sequence (NP_001019389.1) using MultAlin and the BLOSUM62-12-2 matrix [56]. Sequences were considered to be *bona fide* Mo-insertase E-domains if aligning to both, the *E. coli* and gephyrin E-domains. Application of this final control step revealed few sequences to be Mo-insertase G-domain like proteins (tabulated in supplementary table S2).

*Mo-insertase domain classification* – In order to identify E- and G-domain comprising regions in Mo-insertase fusion proteins, the following work flow was carried out: Upon initial identification, obtained sequences were routinely aligned with the G-domain encoding sequence of gephyrin ([22], invertebrate type Mo-insertases), Nit-9 [25], fungal type Mo-insertases) and Cnx1 ([27], plant type Mo-insertases). Due to highest sequence conservation, for *Gnathostome-*type Mo-insertases these have been generally assumed to possess the domain organization identified for mammalian gephyrin [22].

*Visualization of conservation grades* – To visualize the degree of Mo-insertase surface conservation, the degree of amino acid conservation of vertebrate- / invertebrate- and plant-type Mo-insertases (see the results-section for details) was calculated to percent (https://github.com/bpucker/molyb). The conservation scores were subsequently written to the b-factor column of the *A. thaliana* Mo-insertase (PDB code: 6Q32) and the *R. norvegicus* Mo-insertase A-chain (PDB code: 2FU3), respectively. The modified *R. norvergicus* Mo-insertase A-chain has subsequently been duplicated to replace the B-chain in the figures shown by using PyMol [57]. For *A. thaliana* the second monomer was built by crystallographic symmetry. All structure visualization was carried out using PyMol [57].

*3D structural prediction of proteins* – Prediction of monomeric and dimeric Cnx1E models was performed using AlphaFold2 [29] and AlphaFold3 [30]. Modelling with AlphaFold3 was used to generate wt Cnx1E and active site variants S328T, S400T, K297R, K297L, K294R, and R369K. The G296A substitution was predicted to induce backbone rearrangements that could impact functionality (as described in [11]). To reveal any impact, the two deposited conformations of the G296-K297 segment from the Cnx1E crystal structure (PDB ID: 6ETF) were extracted and used as templates for structure prediction of the G296A variant using AlphaFold2 with default parameters [29]. Only the “relaxed” backbone conformation present in 6ETF [11] yielded a viable template for modeling G296A. For structural comparison, Moco-AMP (PDB ID: 6Q32) [12] or molybdate (PDB ID: 6ETF) was superimposed from the solved Cnx1E structures onto the predicted models to visualize substrate/product positioning within the active site.

*Analysis of catalytically important residues* – Vitally important Mo-insertase active site residues were recently summarized for eukaryotic Mo-insertases [6]. The conservation amongst active site residues of various Mo-insertases reported here were checked using KIPEs v3 [28] (https://github.com/bpucker/KIPEs) and the bait input sequences as well as the analyzed residues are listed in Table 3.

*Analysis of other conserved proteins* (MOCS2B and ADH) – The *Homo sapiens* ADH (AAA19002.1) and MOCS2B (NP_004522.1) sequences were searched with BLAST against all downloaded polypeptide sequences using the script collect_best_BLAST_hits.py [51] and default settings. The top hit obtained from the BLAST was used for further analysis. The obtained sequences were subsequently used for the generation of alignments, using MAFFT v7 [52]. Phylogenetic trees were constructed with IQ-TREE2 [53] using the maximum likelihood estimation and 1000 bootstrap replicates. The model finder integrated in IQ-TREE2 identified LG+R8 as the best model. ITOL was used to visualize the phylogenetic trees [54].

## ACKNOWLEDGEMENTS

This work was supported by the de.NBI Cloud within the German Network for Bioinformatics Infrastructure (de.NBI) and ELIXIR-DE (Forschungszentrum Jülich and W-de.NBI-001, W-de.NBI-004, W-de.NBI-008, W-de.NBI-010, W-de.NBI-013, W-de.NBI-014, W-de.NBI-016, W-de.NBI-022).

## DATA AVAILABILITY STATEMENT

The BLAST database is publicly available at NCBI. Also, protein and genome sequences analyzed in this study are publicly available from the NCBI. A summary of all publicly available data, which was analyzed in this study, can be found at https://github.com/bpucker/molyb. Processed data and alignments are available from https://github.com/bpucker/molyb. The surface conservation analysis script can be found on GitHub (https://github.com/bpucker/molyb).

## AUTHOR CONTRIBUTIONS STATEMENT

Tim Julian Schmidt: Acquisition, analysis and interpretation of data Ahmed H. Hassan: Acquisition, analysis and interpretation of data Tobias Kruse: Design of the work, analysis and interpretation of data, drafting the article Boas Pucker: Design of the work, analysis and interpretation of data, revising the article

## ADDITIONAL INFORMATION

### Competing interests

The authors declare no competing interests.

## Notes

https://github.com/bpucker/molyb

### Competing Interest Statement

The authors have declared no competing interest.

