## Supplementary Data for "Phylogenetic based dissection of eukaryotic Mo-insertase functionality: From mechanism to complex assembly"

**S U P P L E M E N T A R Y  M A T E R I A L**

**F I G U R E S**

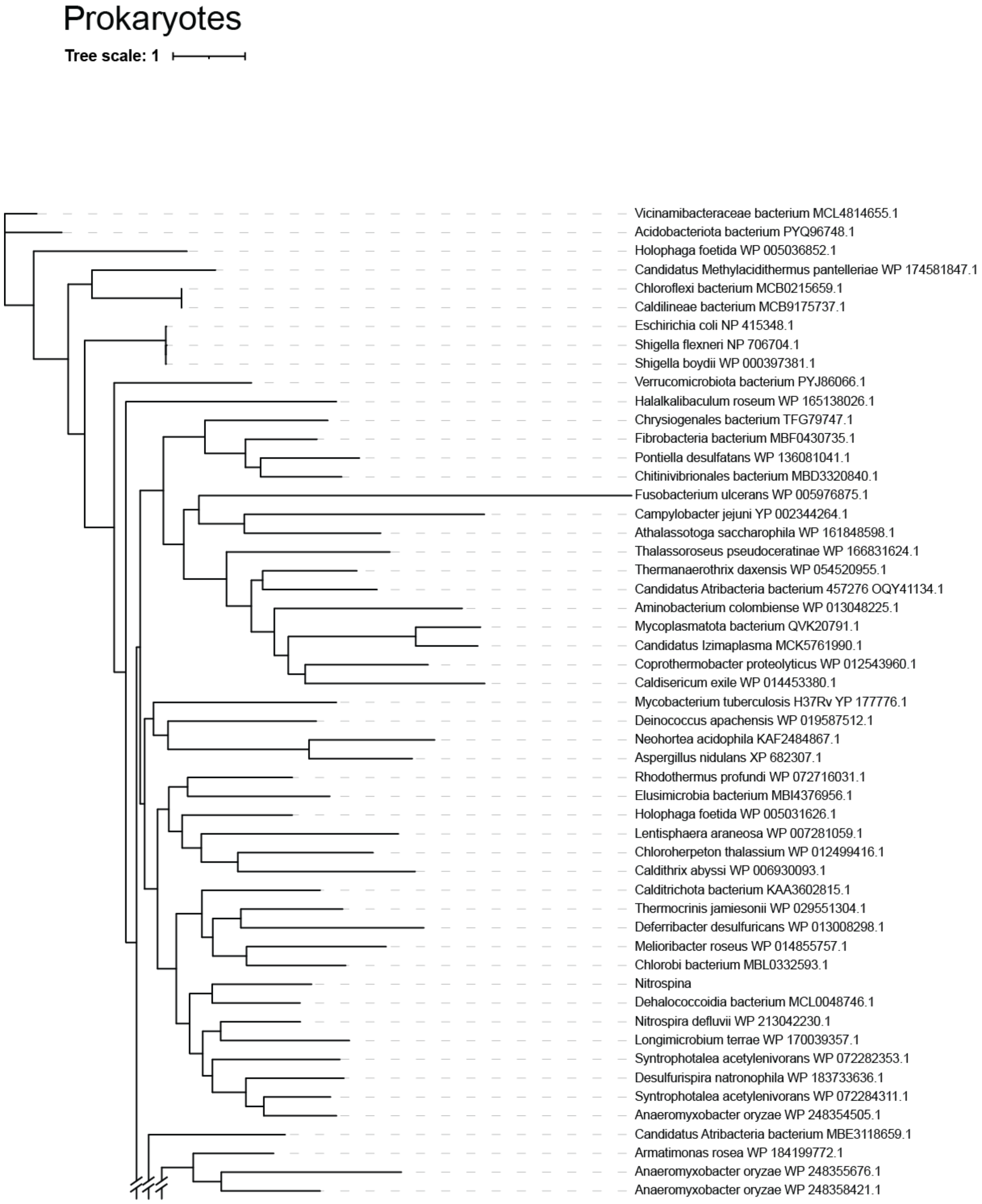

**Figure S1: Partial representation of the phylogenetic distance tree obtained from maximum likelihood: Prokaryotes.** Species name and accession number of the identified MoeA homologous sequence are given next to the branches.

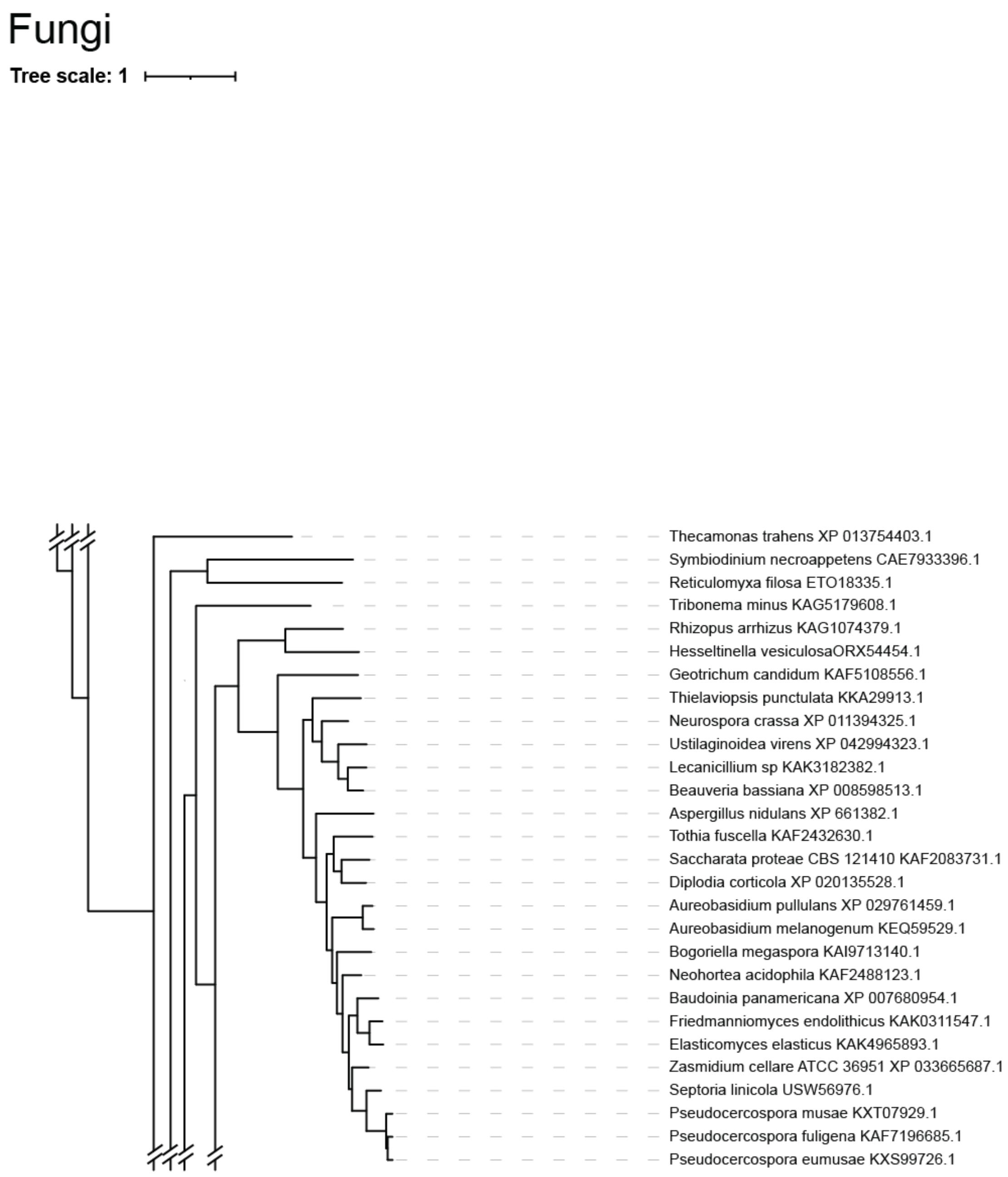

**Figure S2: Partial representation of the phylogenetic distance tree obtained from maximum likelihood: Fungi.** Species name and the accession number of the identified MoeA homologous sequence are given next to the branches.

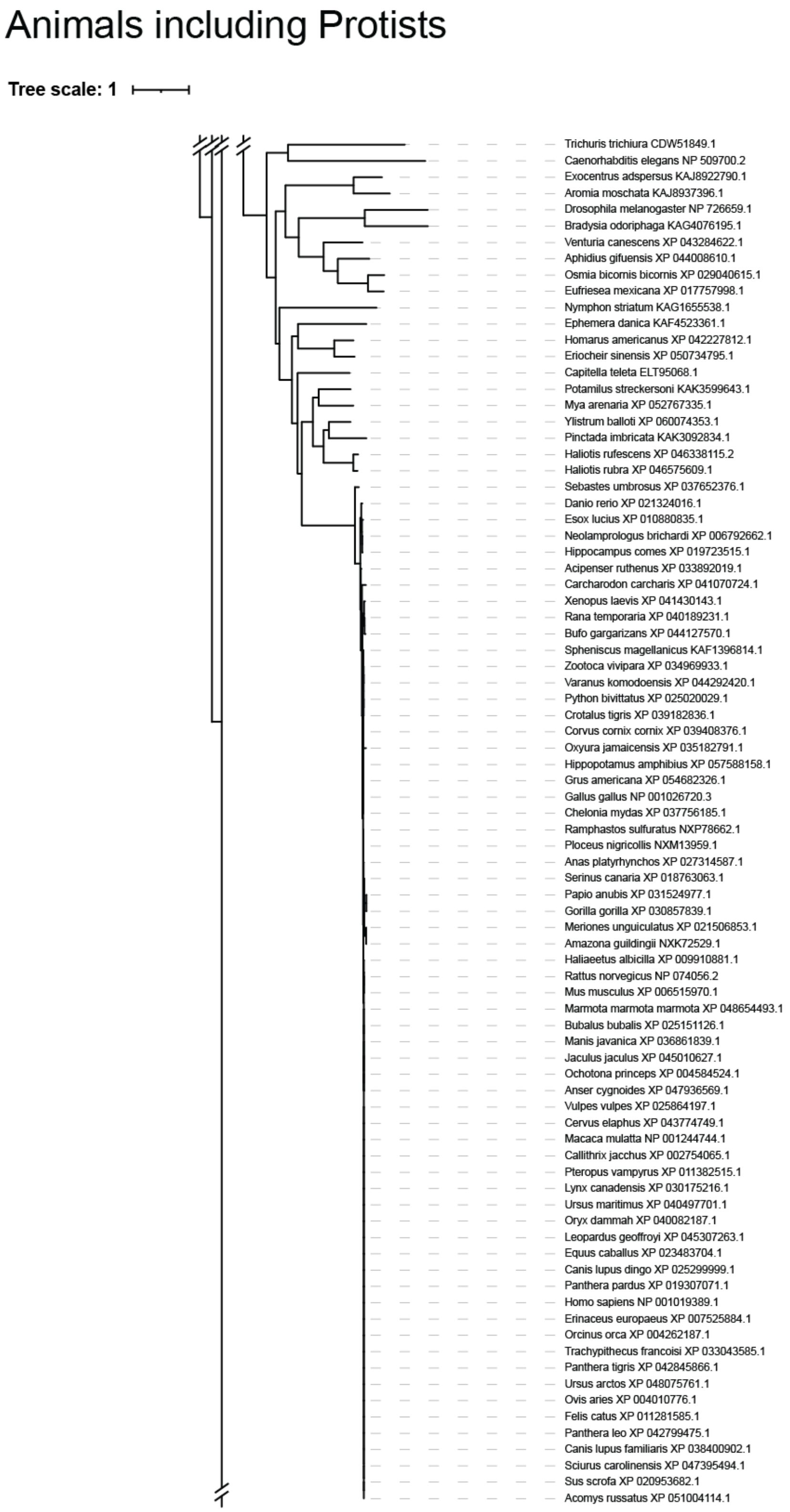

**Figure S3: Partial representation of the phylogenetic distance tree obtained from maximum likelihood: Metazoa.** Species name and the accession number of the identified MoeA homologous sequence are given next to the branches.

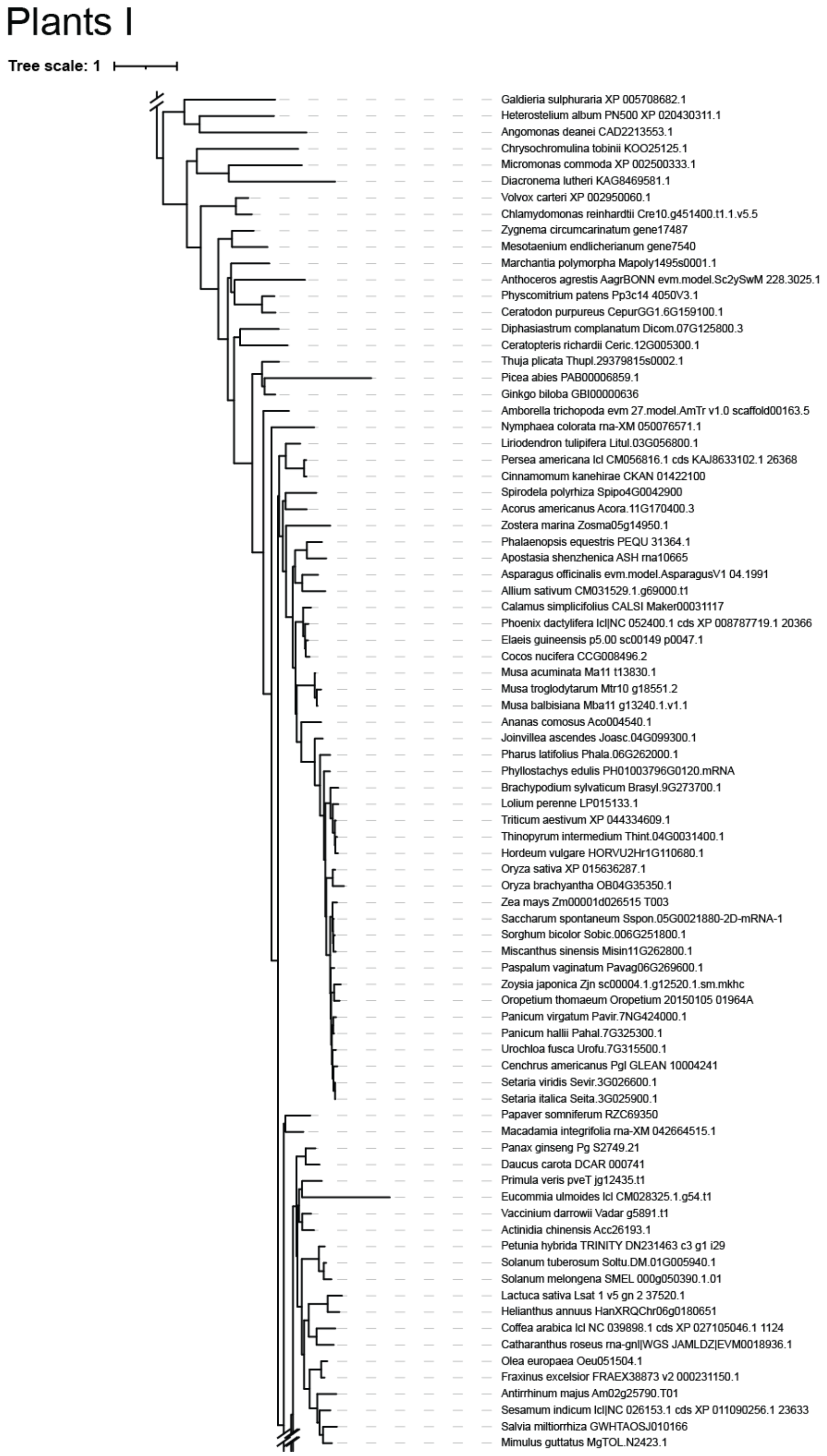

**Figure S4: Partial representation of the phylogenetic distance tree obtained from maximum likelihood: Plants I of II.** Species name and the accession number of the identified MoeA homologous sequence are given next to the branches.

### Plants II

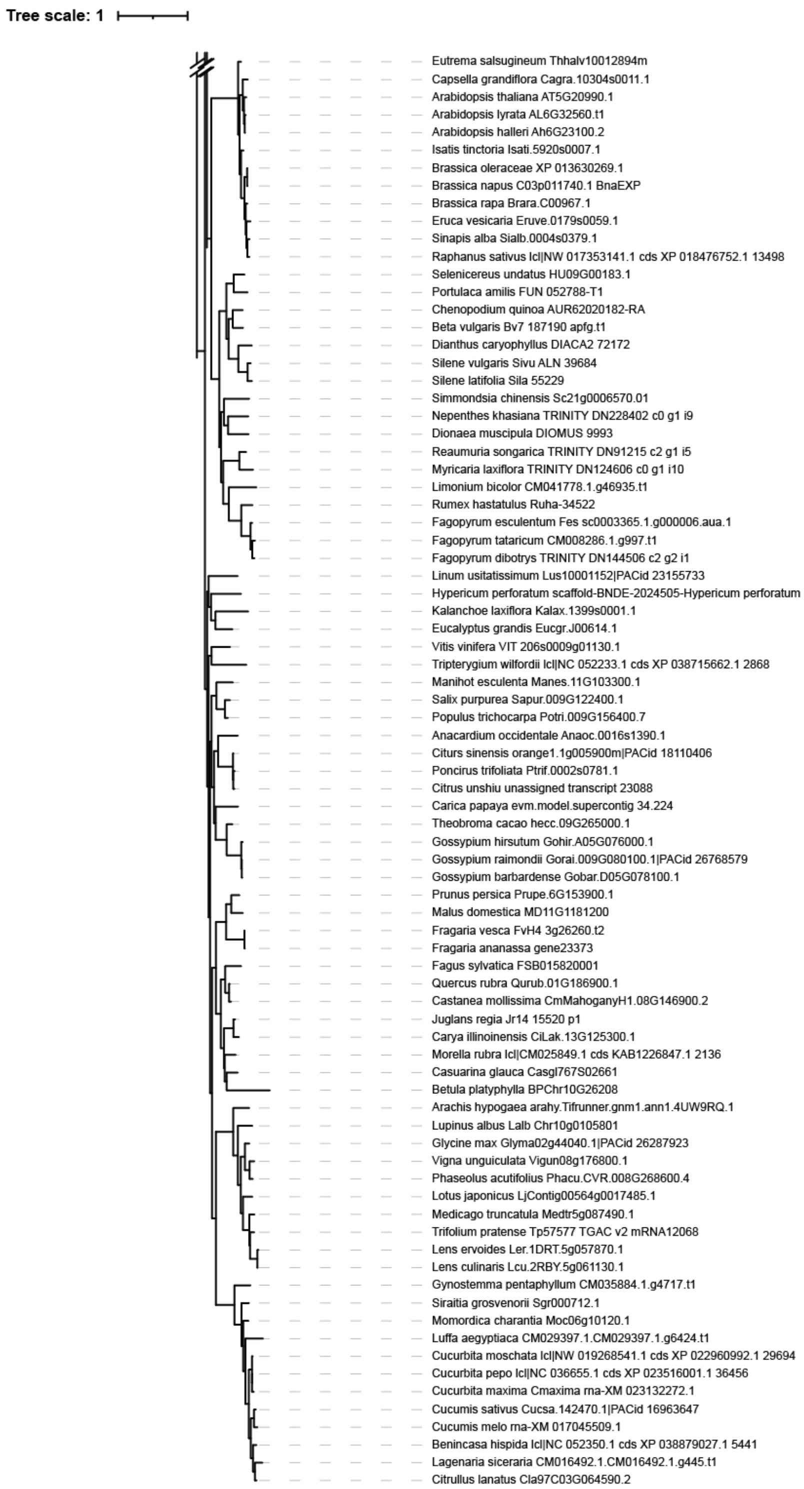

**Figure S5: Partial representation of the phylogenetic distance tree obtained from maximum likelihood: Plants II of II.** Species name and the accession number of the identified MoeA homologous sequence are given next to the branches.

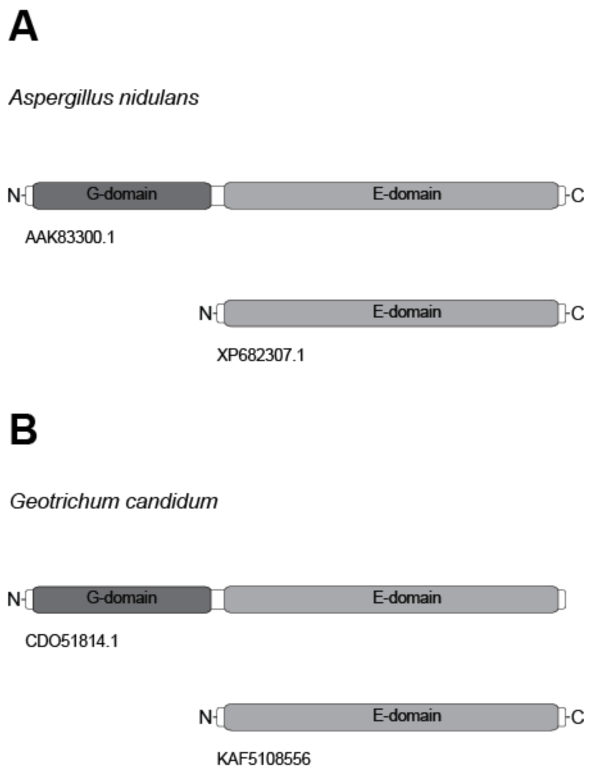

**Figure S6: Schematic representation of identified Mo-insertase domains in *Aspergillus nidulans* and *Geotrichum candidum*.** Mo-insertase domains identified in *A. nidulans* (A) and *G. candidum* (B). (A) and (B): domains were annotated according to *A. nidulans* CNXE domain annotation [1]. Our initial BLASTp approach (see the materials and methods section for details) identified protein XP682307.1 (*A. nidulans*) and KAF5108556 (*G. candidum*). (A) the *A. nidulans* Mo-insertase full length Mo-insertase domain organization is shown according to [1]. The schematic domain organization shown was taken from Fig. 2. (B) the *G. candidum* full length Mo-insertase (CDO51814.1) was identified by an BLASTp search using the NCBI protein database (non-redundant protein sequences (nr), default settings) in the *taxon G. candidum* and using full length *A. nidulans* Mo-insertase (AAK83300.1) as query.

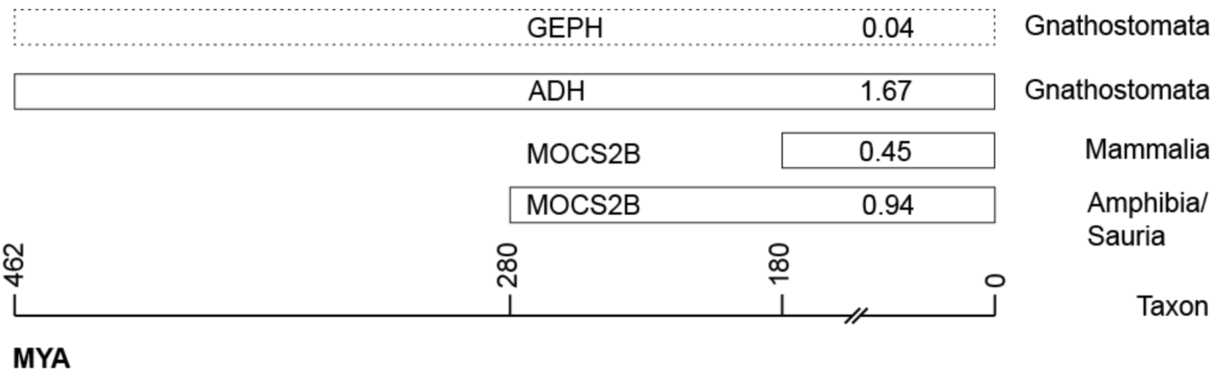

**Figure S7: Patristic distance of the MOCS2B and alcohol dehydrogenase from different taxons.** Patristic distance of MOCS2B and alcohol dehydrogenase from indicated taxons. For comparison the patristic distance determined for gephyrin (GEPH) in the taxon *Gnathostomata* is given. The estimated age when the compared taxons emerged is indicated (MYA = million years ago). For calculation of patristic distances, the taxons *Amphibia* (320 MYA) and *Sauria* (280 MYA) were combined. Estimates of taxon ages were extracted from the evolutionary time tree of life [2, 3].

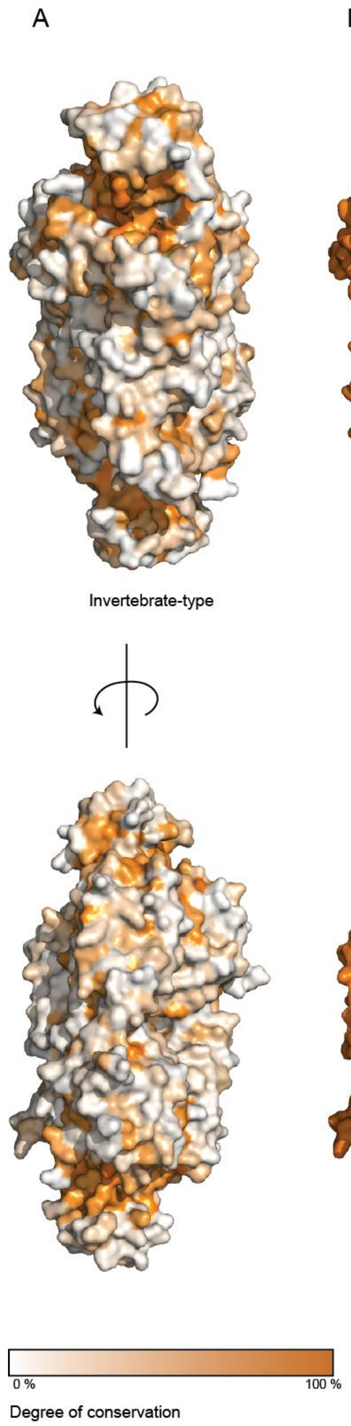

**Figure S8: Conserved residues of eukaryotic Mo-insertases.** Surface representations of the *R. norvegicus* (PDB code: 2FU3, A and B) and *A. thaliana* (PDB code: 6Q32, C) Mo-insertase E-domain. Colors indicate the degree of conservation of surface exposed amino acids amongst members of the Invertebrate-type Mo-insertase (A) the Gnathostome-type Mo-insertase (B) and the Plant-type Mo-insertase (C). The color of the protein surface correlates with the degree of conservation as indicated. The active site as identified for the plant-type Mo-insertase Cnx1[4] is encircled.

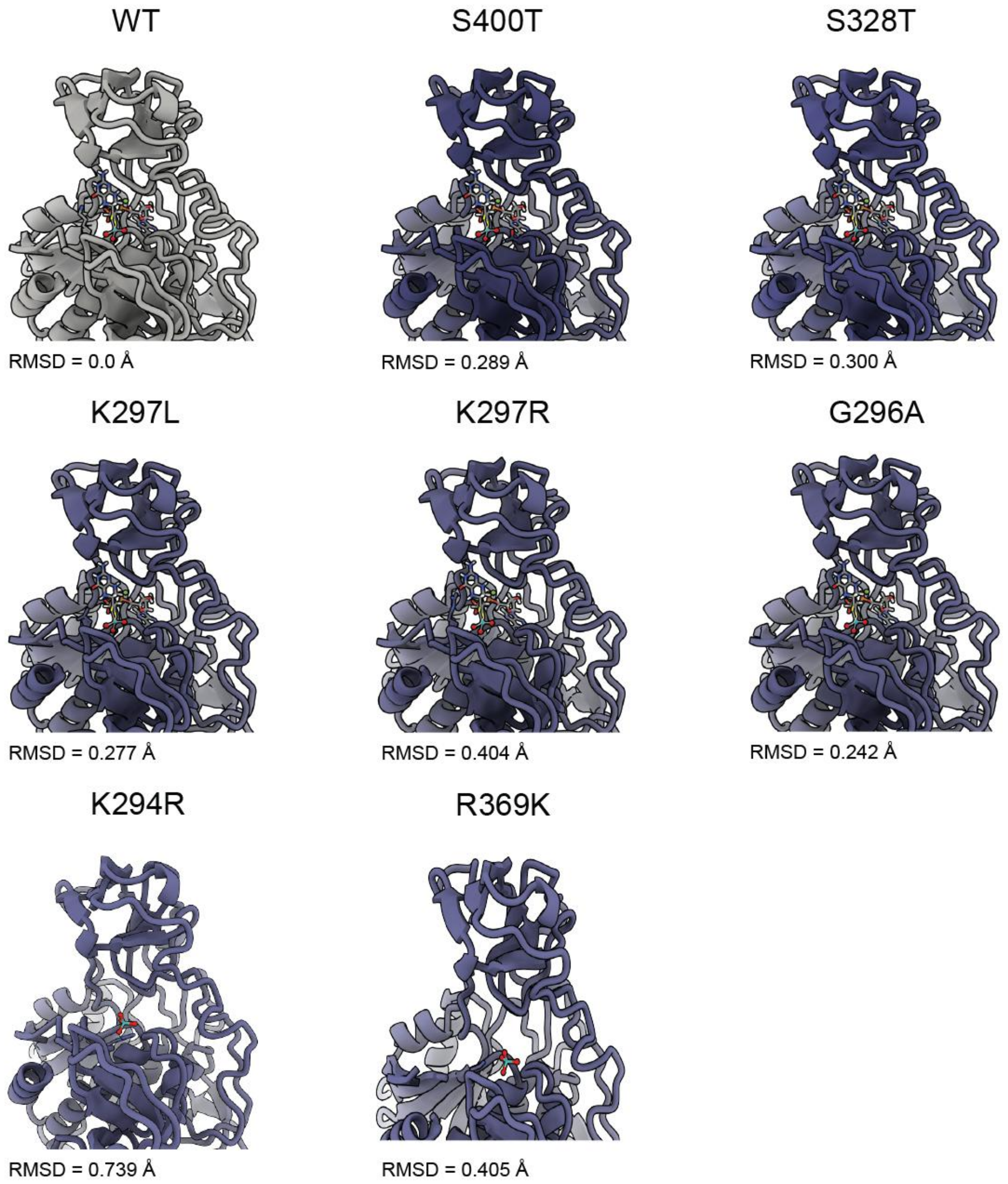

**Figure S9: AlphaFold-Based modelling of Mo-insertase active site variants.** Partial representation of modelled variants (blue) superimposed with the wildtype Cnx1E structure (6ETF, [5], grey). RMSD = root mean square deviation.

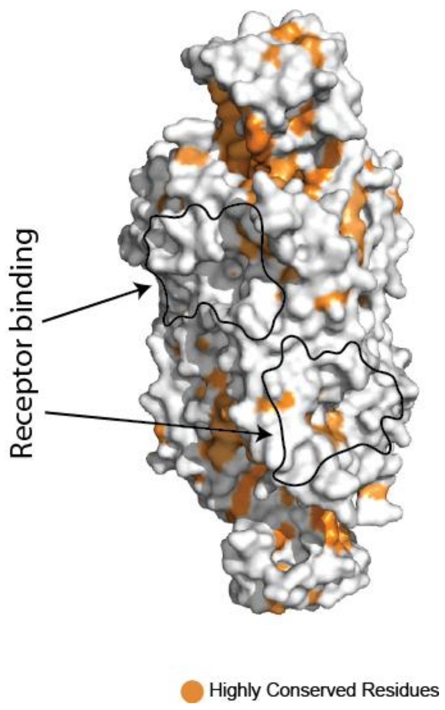

**Figure S10: Highly conserved residues of Invertebrate-type Mo-insertase.** Surface representations of the (A) *R. norvegicus* (PDB code: 2FU3) Mo-insertase E-domain. Left: Highly conserved (> 70% identity) residues of Invertebrate-type Mo-insertases are shown color coded as specified in Fig. 5. residues that fall below this threshold are shown in grey. (A) The plotting refers to invertebrate-type Mo-insertases. The receptor binding site (according to [6]) is encircled.

# T A B L E S

**Table S1: Mo-insertases with a diverging domain arrangement.** Mo-insertases possessing a diverging domain arrangement as compared to the clade (fungi, animals, plants) where these grouped to (see Fig. 2 for comparison) are tabulated. The E-domains tabulated were identified by the initial BLASTp search. If indicated (*i.e.* when a separate existent E-domain was identified, G-domains were identified by using the MogA sequence (QKU47929.1) as query for a BLASTp search and restricted to the respective organism, using the NCBI protein database (non-redundant protein sequences (nr), default settings). As an exception for the identification of the *Volvox carteri* G-domain BLASTp searches (standard settings) were carried out using the JGI database [7] with queries restricted to *Volvox carteri*. If indicated the number of the first and last amino acid of the E- and G-domain within the fusion proteins are given. For *Tribonema minus*, *Symbiodinium necroappetens* and *Heterostelium album* the structure of the G-domain has been predicted using [8] to refine domain annotation. For *Reticulomyxa filose* (ETO18335.1) only a partial sequence was available which showed significant sequence similarities to the *H. sapiens* gephyrin E-domain. The *Pinctada imbricata* G-domain is part of a hypothetical protein comprising 1183 aa. In *Diacronema lutheri*, the E-domain was identified to be part of a hypothetical protein comprising 611 aa (KAG8469581.1) respectively. The G-domains identified in *Micromonas commode*, *Diacronema lutheri* and *Chrysochromulina tobinii* possessed an N-terminal extension comprising ca. 150 residues. Domain classification in fusion proteins was carried out as described in the materials and methods section. The *C. reinhardtii* Mo-insertase domain organization was described elsewhere [9]. For Streptophyta species where separate E-domains were identified, G-domain containing sequences were identified by using the *A. thaliana* Cnx1G sequence [10] as query for a BLASTp searches and restricted to the respective organism, using the NCBI protein database (non-redundant protein sequences (nr), default settings). The *Carya illinoinensis*, *Tripterygium wilfordii* and *Nymphaea colorata* G-domain containing sequences (KAG2674269.1, XP_038715653.1 and XP_031481496.1 respectively) comprises G- and E-domains with the plant type orientation, indicated by an asterisk.

| Clade | Species | E-domain | G-domain |
|---|---|---|---|
| **Fungi** | *Tribonema minus* | KAG5179608.1 | |
| | | (16-448) | (480-620) |
| | *Symbiodinium* | | |
| | *necroappetens* | CAE7933396.1 | |
| | | (10-439) | (468-608) |
| | *Reticulomyxa filose* | ETO18335.1 | |
| | | (partial) | n.a. |
| **Animals** | *Trichuris trichiura* | CDW51849.1 | CDW57105.1 |
| | *Caenorhabditis elegans* | NP_509700.2 | NP_001370857.1 |
| | *Pinctada imbricata* | KAK3092834.1 | KAK3093217.1 |
| | *Ephemera danica* | KAF4523361.1 | n.a. |
| **Plants** | *Carya illinoinensis* | CiLak.13G125300.1 | KAG2674269.1* |
| | *Eucommia ulmoides* | lcl_CM028325.1.lcl | n.a. |
| | | _CM028325.1.g54.t1 | |
| | *Fagus sylvatica* | FSB015820001 | n.a. |
| | *Gynostemma penaphyllum* | CM035884.1. | n.a. |
| | | g4717.t1 | |
| | *Luffa aegyptiaca* | CM029397.1 | n.a. |
| | | g6424.t1 | |
| | *Nymphaea colorata* | XM_050076571.1 | XP_031481496.1* |
| | *Tripterygium wilfordii* | NC_05223.1_cds | XP_038715653.1* |
| | | XP_038715662.1 | |
| | | _2868 | |
| | *Solanum melongena* | SMEL_000g050 | n.a. |
| | | 390.1.01 | |
| | *Micromonas commoda* | XP_002500333.1 | XP_002504805.1 |
| | *Volvox carteri* | XP_002950060 | Vocar.0002s0019.1.p |
| | *Chlamydomonas reinhardtii* | DQ311646.1 | DQ311645.1 |
| | *Galdieria sulphuraria* | XP_005708682.1 | n.a. |
| | *Angomonas deanei* | CAD2213553.1 | n.a. |
| | *Diacronema lutheri* | KAG8469581.1 | KAG8466760.1 |
| | *Chrysochromulina tobinii* | KOO25125.1 | KOO34835.1 |
| | *Heterostelium album* | XP_020430311.1 | |
| | | (245-671) | (11-181) |

**Table S2: Sequences not considered for analysis.** Contaminated sequences and G-domain like sequences identified within the dataset are tabulated. Databases: Pucker *et al.*, 2024 [11]; O'Leary *et al.*, 2024 [12].

| Originally identified in Species / Database | Sequence | Reason for elimination |
|---|---|---|
| *Tamarix ramosissima* / Pucker *et al.*, 2024 | AKPGTMASSTPQSLLRAAILIVSDTASKDPSTDKAAPVLKEV<br>FETEGAGKWDTATAGSSDSNGIAVKIVPDHKAQIQDAITTWA<br>DSEDFYNLIITTGGTGFTPKDNTPEAVSPLLHRHAPGLVHGI<br>LAASFEVTPFAMMSRPVAGVRNKTIIITLPGSPKGAKENLQA<br>VLKLLPHACLQAAGENSRAAHAGGVKQLEKDAGVASFSVKGK<br>TAHAHNRDLGHSHSHDHSHGHGHGGHAVPKAHTSPSQRPPQS<br>NDPRLGASHRARQSPYPMLSVSEAVDTILSHTPSPGPTTAPL<br>SISLVGAVISSDIKAPEAVPAYRASIVDGYAIIVPEDPSQRH<br>TTKGTFPVASVSHAQASSMPPPLQSGEIARITTGAPLPDNAN<br>AVVMVEDTVIASLTSTDPAEEKEVTILTDALVPGENIREPGS<br>DIALNSTILTSGARITGLGGEIGLLAASGTHTVPVYRRPRVG<br>VLSTGDEVTDISHPGPLTGGMIRDSNRPSLLSLISSWRLCSE<br>VVDLGIARDTPPSDLETKLRDAYRVQDLDVVVTTGGVSMGEL<br>DLLKPTIERTLGGTIHFGRVNMKPGKPTTFGSVPVKTNDGTQ<br>RNERLIFALPGNPASALVTANLFLLPALQKLAGIKDVTGLER<br>VDVRLAGRVRCDKSRPEYHRCTVHFDGTSGGLVAVSTGMQRS<br>SRVGSVGGANGLLCLPVKEGHLEEGEKCECLLMGMIVGS | **contaminated sequence**<br><br>**correct assignment:**<br>*Knufia fluminis* |
| *Tamarix ramosissima* / Pucker *et al.*, 2024 | FRKESKWPMIPIDEALQIVLNQTEILPAKKVALKNSLGHIVA<br>EDILAKEPLPPFRASIKDGFAVRFQDGPGVYPVIGTVTAGVI<br>PNFKVESKTIARITTGSALPEGADAIVMVERTELLPDRDEQG<br>RELVRILEGAGEGHDIRMIGSDVAIGELVLKKGERIGPAEVG<br>LLATVGVSDVPVIQSPKIAVLSTGDDLCETDQPLTPGHIRDS<br>NRSMLIAAIRESEVSWNESCID | **contaminated sequence**<br><br>**correct assignment:**<br>*Litorilinea sp.* |
| *Idotea baltica* / O'Leary *et al.*, 2024 | MDCCATQGLIPVETALDTLLSQVSPISNTVVLPLSDAIGFVL<br>ADDICSPINVPPFANSAMDGYAVRISDLEQSLTLPLAGKSFA<br>GIPFDGEWATQTTIRIMTGAKIPTGCDAVIMQELTSDSDGNI<br>TFDLSLTDVKPQTNIRPIGDDVAQGQTVLEKGHRLTPRDIPL<br>IASLGINDIPVVAKPKVAFFSTGDELKPLGQPLEDGQIYDSN<br>RYGIKVLIERFGCEAIDLGIIPDCPETLRKVFLKADKEADVV<br>VTSGGVSVGEADYTKDILDELGQIGFWKLAIKPGKPFAFGNL<br>PNSYFCGLPGNPVSAMLTMYVLVQPMLAKLAGHSSWEAPKSI<br>PAVATTLFKKRPGRTDYQRGIYSINAQGQFEVATTGNQGSGA<br>FSSMSIANCFVVLERERGRVESGEMVNIELFNSSLY | **contaminated sequence**<br><br>**correct assignment:**<br>*Aliivibrio fischeri* |
| *Tamarix ramosissima* / Pucker *et al.*, 2024 | SGGVSMGDRDFVKPLLEEKGKVYFSKVLMKPGKPLTFAEIRA<br>KPTESMLGKTVLAFGLPGNPVSCLVCFNIFVVPTIRQLAGWT<br>SPHPLRVRLRLQEPIKSDPIRPEFHRAIIKWKTTMDQGLPDL<br>LLRALDIR | **contaminated sequence**<br><br>**correct assignment:**<br>*Arabidopsis thaliana* |

| **Originally identified in Species / Database** | **Sequence** | **Reason for elimination** |
|---|---|---|
| *Tamarix ramosissima* / Pucker *et al.*, 2024 | AIQLATGVNTHSHPSNTKSGCRCSDTTINNSKYEMIEYDVAL<br>QTVLDQSSAVTPDTETLLTTESLGRVTASNIHSTVNIPSYNT<br>SYVDGYALSSITTTNTFTVLTSYTAGDTLISSINIGDGECVR<br>VNTGSLIPINTRYIVMVEDTSLSGDTTITLSTAVTAESMEGN<br>IRRIGTDLRIGDLLLHSNHTITPFDIPLLVSAGIHTITCYKQ<br>CSIGILSSGNEVVDISSTNNTTTTTVVDGMIYDTNRPALIA<br>LFKSHGVNIVDFGIVRDERQEIVHRMTESLKQVDVLITTGGV<br>SMGEKDYIKPILEQEMNSTIHFGRVNLKPGKPTTFATLTLPD<br>DKKKKKFVFGLPGNPVSAIVTSILFVLPLARKMSGHAGYLNE<br>AIPVVIGNDIHLDSRREFMRATVTKEVIGNGGSVRYVATSCG<br>KQTSSRLLSMKGADVLLKLPPREKVGGDGILKKGSVVDAILL<br>K | **contaminated sequence**<br><br>**correct assignment:** *Basidiobolus meristosporus CBS 931.73* |
| *Cullicoides impunctatus* / O'Leary *et al.*, 2024 | MEPFTQGLIALDDALKVMLESITPLTDDELSMLREASGRITA<br>TAIISPVDVPPFANAAMDGYALRYADFATDRIFPVAGKALAG<br>FPFSEPWPTGSCVRIMTGAPLPAGADVVIMQELATVEGDGVR<br>FNQPVSPGQHIRLAGEDICAGSAIIEPGRRLGTAQLPLCASL<br>GLNQLSVIRRPKVALFSTGDELQLPGQPLSEGQIYDTNRLAV<br>GIMLEKLGCEVRDFGIIPDCPETLKHTFTEADSWADVTISSG<br>GVSVGEADYTRTILESLGKITFWKLAIKPGKPFAFGRLQHSW<br>FCGLPGNPVSAVVSFYQLVQPLLRHLAGEKNITPPRLRARLD<br>GRIKRNPGRIDFQRGRLYQSPEGQFRVTTTGAQGSHVFSSFN<br>QANCFIVLPREQGDVSPDEWVEAEPFNHLLQG | **contaminated sequence**<br><br>**correct assignment:** *Candidatus Erwinia impunctatus* |
| *Ephemera Danica* / O'Leary *et al.*, 2024 | MTDNSSCISCATENNLLSVAEAREHMIAEVQSITGREFLSLR<br>NALGRVLATDIIAPHDVPAHDNSAMDGYAVCFDSLAAEGETR<br>LTVVGTAFAGNAFSGQVGRGQAVRIMTGAVLPAGADTVVVQE<br>VVRREGSEVVVPAGQQVGQNTRRAGEDLARGAVALPAGKRIG<br>PAELGLVASLGVAEVAVKRRLRVAFFSTGDELASIGKPLAPG<br>EVYDSNRYTLHGLLTRLGAEIIDLGVVPDRPEALEAALAEAA<br>QIADAIITTGGVSVGEADFVREILDKLGEVRFWKINIKPGRP<br>MAFGKVGKAWLFGLPGNPVAVMVSYTQLALGALYRLSGLDPL<br>PERPLLAAISANPIRKQAGRREYLRGRIAAVDGAWQVKTAGN<br>QGSGVLRSMSEANCFVVLPEDCTSVASGDPVAVELFDGLF | **contaminated sequence**<br><br>**correct assignment:** *Dechloromonas sp.* |
| *Trichuris trichiura* / O'Leary *et al.*, 2024 | MEFTTGLMSLDTALNEMLSRVTPLTPQETLPLVQCFGRILAS<br>DVVSPLDVPGFDNSAMDGYAVRLADIASGQPLPVVGKSFAGQ<br>PYHGEWPAGTCIRIMTGAPVPEGCEAVVMQEQTEQTDNGVRF<br>TAEVRSGQNIRRRGEDISAGAVVFPAGTRLTTAELPVIASLG<br>IAEVPVIRKVRVALFSTGDELQLPGQPLGDGQIYDTNRLAVH<br>LMLEQLGCEVINLGIIRDDPHALRAAFIEADSQADVVISSGG<br>VSVGEADYTKTILEELGEIAFWKLAIKPGKPFAFGKLSNSWF<br>CGLPGNPVSATLTFYQLVQPLLAKLSGNTASGLPARQRVRTA<br>SRLKKTPGRLDFQRGVLHRNADGELEVTTTGHQGSHIFSSFS<br>LGNCFIVLERDRGNVEVGEWVEVEPFNALFGGL | **contaminated sequence**<br><br>**correct assignment:** *Escherichia coli* |
| *Russula earlei* / O'Leary *et al.*, 2024 | MITVNEAKNIIRHNCKVLPAITLPLESALTYVLAEDVYAVAD<br>IPAFDQSSMDGYAIAFDDYYYHHKLQVEGVIPAGHSIAARIQ<br>ARQAARIFTGAPMPQGADTVIIQEKVTVENNELTGTDTLLKK<br>GSNVRPKGSEIKAGEIAATKGTCLSPAAIGFLAGIGIAEVKV<br>ISKPTISIVVTGNELRQPGKPLLHGQVYESNSFTLNAILQQL<br>FMSNVTTIWVDDDLCLMEDALEKALEHSDMVLLTGGVSVGDY<br>DYVLEAASICGVQQLFHRIKQKPGKPLFFGKKGDKLVFGLPG<br>NPSSVLTCFYEYVIPALQQLTQRKSIIKVVHLPLAKAHQKKP<br>GLTHFLKGHVEANKVIPLQAQESYRLSSFSAANCLIRLEEEG<br>EEYAAGAMVEVHLLPF | **contaminated sequence**<br><br>**correct assignment:** *Flavisolibacter sp.* |

| Originally identified in Species / Database | Sequence | Reason for elimination |
|---|---|---|
| *Symbiodinium necroappetens* / O'Leary *et al.*, 2024 | AFLVGESLMRQAEVTAATRALLAGKAVMVDVGAKAETERRAT<br>ARGEVVMQPETLRLIESGGVQKGDVLSVARLAGIMGAKRTPE<br>LIPLCHPLALTSVTLDLRLRPERDAVEIEATCKLTGRTGVEM<br>EALTAVSVAALTVYDMCKAVDRGMRIDNVRLVHKTGGKSGTY<br>EDDEAECFFATARPRDARPGDREPPQRSSGMISVEEAQTRVL<br>AAFSPLPAETVAVNQALGRVLAEDVTARVTQPPADVSAMDGY<br>AVRAEDLAEIPARLQVVGRVPAGGRYADRLAPGQAVRIFTGA<br>PLPDGADTIVIQEDCTAEGDAVVVRAGAARGTYVRPAGLDFR<br>AGSLGVPAGRPVSVRDLGLIAAMNRPWVSVRRRPRVAVLATG<br>DEVVMPGDPLGPSQIVSSNGLALCALVEACGGSAINLGIAAD<br>SAESLQRLAAGAAGADLLVTTGGASVGEHDLIRSVLGEAGLE<br>LDFWKIAMRPGKPLMFGRLKDTPMLGLPGNPVSSLVCGLLFL<br>RPMLDRLLGLQRPAHPLEPALLGADLGANDRRQDYLRASLET<br>DADGRRVATPFGRQDSSMLATLTQADALIVRPPHAPALAAGQ<br>VVALLRFPAGLGSIDVRQATRWEAAGPLRASNKGPDDMLTRK<br>QHELLLFLHAHLGEHGVSPSFDEMKEALGLKSKSGIHRLITG<br>LEERGFIRRLPHRARAIEVLRLPEDMAGKSGFAPNVIEGGRR<br>AGLAGARVATDSEAVSLPLYGRIAAGTPIEALRDHSNYVDVP<br>ADLLSRGEHYALQVEGDSMVEAGILDGDTVVIERSDQAENGA<br>IVVALVDDAEVTLKRFRRRGGAIALEPANRNYEPRLFPPDRV<br>KVQGRLIGLLRRY | **contaminated sequence**<br><br>**correct assignment:** *Kiloniellales bacterium* |
| *Tamarix ramosissima* / Pucker *et al.*, 2024 | DYLKQVLDIDLHAQIHFGRVFMKPGLPTTFATLDIDGVRKII<br>FALPGNPVSAVVTCNLFVVPALRKMQGILDPRPTIIKARLSC<br>DVKLDPRPEYHRCILTWHHQEPLPWAQSTGNQMSSRLMSMRS<br>ANGLLMLPPKTEQYVELHKGEVVDVMVIGWL | **contaminated sequence**<br><br>**correct assignment:** *Myotis davidii* |
| *Persea americana* / O'Leary *et al.*, 2024 | MMARPLAGVRHNTLVVTLPGSPKGAVENLQAIIKLLPHACQQ<br>AAGSNSRTLHAGGVAQLEKDAGVSSGSHSHNHDHKHHHGHSH<br>GHSDKHTHGHAVPRAHTTAAERLASNDPTAGPTRRYRESPYP<br>MLSVKDALDVIANNTPKPVAYRRPVDEDLVGHVLAEDVPAKE<br>SVPAFRASIVDGYAIIASKHVMVPSTKGIFPVVSISHAKAGS<br>VEKLEIGEVARITTGAPLPPGATSVVMVEDTVLRKSTDDGKE<br>EAEIEILTDAIEPGENVREVGSDVTAGDIILRKGEGVSATGG<br>ELGLLASVGTDTVLAYRKPRVGVLSTGDEIVPHNRQGALQLG<br>EVRDTNRPTLLTSIRAQGFDAVDLGIASDAPGALETTLRNAM<br>REVDVIVTSGGVSMGELDLLKPTIERQLGGTIHFGRVSMKPG<br>KPTTFATIPFKENDGQDTKRLIFSLPGNPASAVVTFNLFVLP<br>ALHQHSGVEPAGLPKIKVVLEQDVRCDEKRDEYHRVVIIAKG<br>DGRLYASSTGGQRSSRIGSFKSANGLLCLPAKNGSIKKGEVC<br>DALLMARLLGEA | **contaminated sequence**<br><br>**correct assignment:** *Peltaster fructicola* |
| *Anaerolineae bacterium* / O'Leary *et al.*, 2024 | MKHSPNSPILLNSSLHVDEARKAITNLVSELQQESSILNDPA<br>DIETVSLDHAINRVLAQDLLSPIDVPAADNSAMDGYAFDGKC<br>LSQAGSEVTLNIVGTALAGKPFEGEIGQGECLKIMTGALMPA<br>DCDTVIPQEFTTSASAESICFPSNQLKAGENRRLRGEDLQKD<br>KAAISAGRLLRPSDLGLAASLGTSHLQVRRKLRVAILSSGDE<br>LRSLGQPLDPGSIYDSNRYSLNGMLNRLNIDIIDCGIVRDNP<br>DSLKDAFIAAASKADVLISSGGVSVGEADFTKQVMLELGDVG<br>FWKIAMRPGRPMAFGTLKPVPSKSPARKTLFFGLPGNPVAVM<br>VTFYQFVRSALLQLGGVTQADLPLVQAISENAIRKKPGRTEF<br>QRAILGRNVDGKPSVRITGSQGAGILRSMSEANCFVILRHDQ<br>GNVAPGELVDIALFEGLL | **contaminated sequence**<br><br>**correct assignment:** *Polynucleobacter sp.* |
| *Tamarix ramosissima* / Pucker *et al.*, 2024 | RAMLLSAAVQQNCKIIDLGIARDDEEELERIFNKAFDAGVDI<br>ILTSGGVSMGDRDFVKPLLEKRGKVYFSKVCMKPGKPITFAE<br>FNLKPAEDTSANKVLAFGLPGNPVSCLVCFHLFVVPAIRHLS<br>GWANAHSLRVQARLQWPIRADPVRPEFHRAIIKWKLNDGSGI<br>PGFVAESTGHQMSSRLLSMKSANALLELPATGSVIPAGTSLA<br>AILISDLSGTPASENSLSDDKVFSIQECVKPTATDVLSTTSV<br>RVAILTVSDTVASGTGPDRR | **contaminated sequence**<br><br>**correct assignment:** *Populus deltoides* |

| **Originally identified in Species / Database** | **Sequence** | **Reason for elimination** |
|---|---|---|
| *Tamarix ramosissima* / Pucker *et al.*, 2024 | LVRHDAERALHEGEIRDSNRPALISCLMAWGIETVDLGIVSD<br>SSKDLETVLRDALRGTMDHPPVDVIITTGGVSMGERDLLKPT<br>IERLLGGTIHFGRVAMKPGKPTTFATIPRKTSNNPRKQVAIF<br>ALPGNPASALVTMHLFVLPALHKLMGFSHPDGTEDRPSRGLP<br>RVRAVLAHPIPRDPKRTEYHRAVVTASRDGRLRASSTGLEGV<br>GQRSSRVASMAKANALLVLPPGVESLPEGELVEALMMGQIVP<br>GN | **contaminated sequence**<br><br>**correct assignment:** *Thermomyces dupontii* |
| *Nosema bombycis* / O'Leary *et al.*, 2024 | MQEQAEQTDEGIRFLAPVKNGQNIRRLGEDIAHGAVVFPAGT<br>RLTAAELPVIASLGIAEVEVVRKVRVAVFSTGDELQLPGQPL<br>ADGQIYDTNRLAVHLMLQELGCEVINLGIIPDDPAKLREAFI<br>QADQQADVVISSGGVSVGEADYTKAILEELGEIGFWKLAIKP<br>GKPFAFGKLNHSWFCGLPGNPVSATLTFYQLVQPLLAKLSGN<br>VGQAQPMRLRVRAASGLKKSPGRLDFQRGVLQRGPDGELVVS<br>STGHQGSHIFSSFSLGNCFIVLERERGNVEAGEWVEVEPFNH<br>LFGGL | **contaminated sequence**<br><br>**correct assignment:** *Klebsiella michiganensis* |
| *Apophysomyces sp. BC1034* / O'Leary *et al.*, 2024 | MSTPNDTAPVACTSHAAPGASTDVPRPPPLSTHEALARVLAA<br>ALPLCAQPEAIEQVPTLDALNRVLAADIRSALDVPPADISAM<br>DGYAVRAADVVAGRPMRVSQRIPAGHPAAGVLEAGTAARIFT<br>GAPLPAGADTVVMQEQARVQSDTVVFDAAPPAGDWINRRGSD<br>IGKDAVILPAGTRLTPQALGLAASVGCAVLPVARRPRVAIFF<br>TGDELTMPGEPLREGAIYNSSRFTLGSLLAALGCDVTDLGIV<br>PDRFDATRDALRRAALEHDLILTSGGVSVGDEDHVRAAVQAE<br>GTLDRWQIAMKPGKPLAFGTVRRALGGESHDADTGTARDTAF<br>FIGLPGNPVSSFVTFVLFVRPFILRLAGAARVEPQALRMRAD<br>FTQKKADRRNEFLRARINDGGGLDLFPNQSSAVLTSTVWGDG<br>LIDNPPGHPIQAGETVRFLPFSALLAVREAVGTAAETVDVPD<br>GVATLGDVRTWLRSRGGAWADALADTRALRMACDHVMTGPST<br>QLTDGCEVAFFPPVTGG | **contaminated sequence**<br><br>**correct assignment:** *Mycetohabitans sp. B8* |
| *Apophysomyces sp. BC1034* / O'Leary *et al.*, 2024 | MTSLQTIASCIADYDPNALPVSAARAIVRQWATPVATVERLA<br>LREALDRVLADDVVSPLDVPAHDNSAMDGYAFDGTALERGGT<br>IRLRVAGTALAGRPHDARVRTGDCIRVMTGAMLPPDCDTVVP<br>QEQVEIDVNGSIQFPATALTRGANRRRAGEDLRAGHAALLAG<br>RTIRASDLGLLASLGIAEVPVRRRLRAAFFSTGDELRSLGQP<br>LEPGCVYDSNRYTLYGMLRRLNLDVIDLGVVPDNRRALETTL<br>RNAAATADVVLSSGGVSVGDADYTRELMDTLGDVAFWRVAMR<br>PGRPFAFGRIGSGAHASESRDALYFGLPGNPVAAMVAFYQIV<br>RDALIAMTGALPHPAPLVRASALDAIGKRPGRTEYPRGIARR<br>RDDGHWEVTLTGAQGSGILRSMSDANCFVILDHHRGPVAVGD<br>TGRAATFANARRRFDMKKQISYIAPGQTAKALILVYLTFSVP<br>IMLLGVLVALIRYGSVELSTVFSALILNALLGFVLLWIACRA<br>YNWVASRFGGIEIVLSDASEEA | **contaminated sequence**<br><br>**correct assignment:** *Mycetohabitans* |
| *Astraeus_odoratus_KAG6335354.1* / O'Leary *et al.*, 2024 | MGAFVVTPQEPTMDFTAGLMPLETALSQMLDRISPLHDVETL<br>PLVRCFGRIAARDIVSPMNVPGFDNSAMDGYAVRLADLQTGN<br>ALPVAGKAFAGQPFGSEWPAGTCVRIMTGAPVPQGCDAVVMQ<br>EETEQTDDGVRFTANVKAGQNIRRTGEDITLGATVFAAGQKL<br>TVGELPVLASLGIAEVDVVRKVRVAVFSTGDELQLPGQPLQD<br>GQIYDTNRLAVHLMLEQLGCEVINLGIIPDDPEKLRAAFIEA<br>DKSADVVISSGGVSVGEADYTKTLLEELGEIAFWKLAIKPGK<br>PFAFGKLPHSWFCGLPGNPVSAALTFYQLAIPLLAKLSGNKA<br>SPLPERLRVRAATRLKKSPGRLDFQRGILARNADAPSARATA<br>LSCWSVSAATWKPANGLRLSALTTCSEADMTVELSDQEMMRY<br>NRQIVLRGFDFEGQEALKAAKVLVVGLGGLGCAAAQYLAAAG<br>VGRMTLLDFDTVSVSNLQRQTLHSDATVGQPKVDSARTALAR<br>INPNVQFTLIDAMLDDDALFAQIAQHDLVLDCTDNVAVRNQL<br>NAGCFAHKTPLVSGAAIRMEGQISVFTYADGEPCYRCLSRLF<br>GENALTCVEAGVMAPLVGVIGSLQAMEAIKVLAHYGTPAAGK<br>IVMYDAMTCQFREMKLMRNPGCEVCGV | **contaminated sequence**<br><br>**correct assignment:** *Enterobacter* |

| **Originally identified in Species / Database** | **Sequence** | **Reason for elimination** |
|---|---|---|
| *Capitella_teleta_ELT99908.1* / O'Leary *et al.*, 2024 | GNNIRKAGEDILEGSQVFTPGRKVRPQDIGLLASLGIANVTV<br>YQKVKVAVFSTGDELKLPGEPLRHGDIYDSNRFVIKAMLKKM<br>EIDIIDLGKIPDDKEQLRQAFLRADREADAVISSGGVSVGDA<br>DYTKEILDELGETGFWKLAIKPGKPFAFGQLPNSVFFGLPGN<br>PVSATVTFQQLAAPALRHMMNQQAEEKVELTLATKTRLKKRP<br>GRRDFQRGKLVYSNSGELQVLSTGNQGSGVMSTMSQSDCYIV<br>LAEEDGDKQAGDLVKVQLFDELLK | **contaminated sequence**<br><br>**correct assignment:** *Endozoicomonas atrinae* |
| *Rhodothermus_profundi_WP_072714208.1* / O'Leary *et al.*, 2024 | MKIILLTIGDELLTGTTVNTNAAWLGAELTAHGFTVVRAETL<br>RDDPHAICQALQRARAEADVVIVCGGLGPTHDDRTREALADC<br>LNRPLQMHAALLEQIKAYFTRRGRHMPERNQVQALVPEGFTP<br>IPNPLGTAPGLWLQDASGIVVVLPGVPHELQGLMREAVLPRL<br>QQLPGRPAILQRTLVTAGIGESELQQRLQGVESLLDEAVQLA<br>YLPSPYGVRLRLTARAASREAAQQRLSELVAFIQKRIQPYLV<br>SLSGETLEVVVGKLLRRLGATVAVAESCTGGHLADCITNVSG<br>ASTYFRGGVVAYDNAVKVEVLGVAPELLAREGAVSEAVAIQM<br>ARGVRKRLGAQVALATTGIAGPTGGTPDKPVGTVWIGLADDQ<br>VAFAQRYFLPDDRKRFKQRATAAALDLLRRHLLQQKVPRAAI<br>FHR | Similarity to G-domain |
| *Anaeromyxobacter_oryzae_WP_248360219.1* / O'Leary *et al.*, 2024 | MIVEILSTGDELLTGQVVDTNSTWLMDRLWDLGVMVRRKTLV<br>ADDRADLVAAIRETAARAELVVMSGGMGPTEDDLTAECVAAV<br>LGVPLELHESSLRVIEERFRKFGRTMTPNNRKQAMFPRGAEV<br>IPNRFGTAPGFAVKVARGEVVCLPGVPLEFKGLADEWVLPRL<br>AARLGDVPAARVLKLFGVPESHADDAMRPVMDDPANAGVRWG<br>YRAHWPEVHVKWTVPGPDAAARADRIRDAVRAIFGEAVWGEA<br>KEELPELVVARLAARGERVALAESCTGGLLAELVTRVPGASN<br>VIDLGVVAYANAMKERLLGVPAGVLAVEGAVSEPVARALAEG<br>ARRVGGAAWGVGITGIAGPTGGTPEKPVGTVHVALASAAGTV<br>HVERQYRGDRERIRRQAAYEALNLLRLALR | Similarity to G-domain |
| *Holophaga_foetida_WP_005034889.1* / O'Leary *et al.*, 2024 | MRIECIAVGSELLSTGRLDTNSVWITERLGRLGLSIHRKTAI<br>GDEPGDLRALFLEAIQRSELVICTGGLGPTFDDLTKETWAEV<br>FGAELVEDPQVRRDILDFYAARNRVPPESNFKQALVPVGARI<br>LRNPFGTAPALYWESPQGYPGRRVILLPGVPLEMKQIWEGQI<br>EALLAPLAQASVHTLRMVVGSVPESTLDERTRALREQHGALE<br>WTILAGISHVELVARGADPALLEAARKGFEGELGEDLVCVGE<br>GSIESTVLDLLQARGETLGLAESVTGGFIATRLVAVPGASQA<br>FLGDVVTYSARAKVQLAGVPERVIQVHGTVSEATTRAMAEGI<br>RDRLGATWGLATTGNAGPSQDAQGPAAVGTIHIALAGPAGTQ<br>TIQYSLPGFRSDIQSRAAAWAMDFLRRRLV | Similarity to G-domain |
| *Nitrospina_gracillis_WP_042252643.1* / O'Leary *et al.*, 2024 | MKNKYDIPQAEIVAVGNELLNGLVSDTNSTFICGQLRMHGLQ<br>VGRISVVGDDADAIRSALDQALSRVDLVIVTGGLGATHDDIT<br>KDVLADYFGTPLVRDPKVEEMIRVFFEKRQRPVPDAALRQAE<br>VPKDGRPLYNDQGTAPGLMFERGEQRVYVLPGVPREAEHLTR<br>QYILPDVAPAGNLCLQQRMLWTTGLVESALWEMFGPVDELEN<br>LVQVASLPSHLGVRIHLTAYGENVEETSAKIEQAETLLEKVL<br>SSYIYARDEQTMESVLGQLLVDRGETVAVAESCTGGLIGHRL<br>TNIPGSSRYFLQGWLTYSNEAKVKSLGVDAALVERHGAVSEE<br>VARAMAEGARQCAGTDWAVSVTGIAGPDGGTATKPVGLTYIA<br>VAGKTLTSCQKFVFPQDRLRNKERAAQAALNLLRLHLIGLK | Similarity to G-domain |

## D A T A  F I L E S

**Data file S1:** Species names and sources of the sequence data sets per species used for BLAST-based analyses for the discovery of MoeA orthologs (xlsx format).

**Data file S2:** Pairwise sequence alignments of all identified fungal Mo-insertases with the *N. crassa* E- and G-domain sequences [1]. The sequence alignments were carried out using EMBOSS Needle Pairwise Sequence Alignment [13] and standard settings. We identified KAG6331869.1 from *Astraeus odoratus* to possesses the fungal domain organization, however the fusion protein was found to be part of a larger protein sized 2058 residues. For XP_007680954.1 from *Baudoinia panamericana* we confirmed the fungal type domain organization, however the G-domain was identified to be truncated.

**Data file S3:** Pairwise sequence alignments of all identified invertebrate Mo-insertases with the *R. norvegicus* E- and G-domain sequences [14]. The sequence alignments were carried out using EMBOSS Needle Pairwise Sequence Alignment [13] and standard settings. We identified KAJ8937396.1 from *Aromia moschata* to possess a N-terminal fused, partial G-domain sequence. For XP_046338115.2 from *Haliotis rufescens*, structures of E- and G-domain were predicted using [8] to refine domain annotation. According to this, the G-domain spans residues 13-175 and the E-domain residues 302-719. For CDW51849.1 from *Trichuris trichiura*, NP_509700.2 from *Caenorhabditis elegans*, KAK3092834.1 from *Pinctada imbricata* and kAF4523361.1 from *Ephemera danica*, we identified no G-domain encoding part within the annotated sequence (see supplementary table S1).

**Data file S4:** Pairwise sequence alignments of all identified plant type Mo-insertases with the *A. thaliana* E- and G-domain sequences [10]. The sequence alignments were carried out using EMBOSS Needle Pairwise Sequence Alignment [13] and standard settings. For the Streptophyta sequences CiLak.13G125300.1 from *Carya illinoinensis,* FSB015820001 from *Fagus sylvatica*, CM035884.1.g4717.t1 from *Gynostemma pentaphyllum*, CM029397.1.g6424.t1 from *Luffa aegyptiaca*, XM_050076571.1 from *Nymphaea colorata*, NC_052233.1_cds_XP_038715662.1_2868 from *Tripterygium wilfordii* and SMEL_000g050390.1.01 from *Solanum melongena* we identified no G-domain encoding part within the annotated sequence (see supplementary table S1). For FvH4_3g26260.t4 from *Fragaria vesca* a C-terminal fused, partial G-domain sequence was identified. For the *Chlorophyta* sequences XP_002500333.1 from *Micromonas commoda*, XP_002950060 from *Volvox carteri* and DQ311646.1 from *Chlamydomonas reinhardtii* likewise no G-domain encoding part within the annotated sequence was identified (see supplementary table S1). For (lcl_CM028325.1.lcl_CM028325.1.g54.t1) from *Eucommia ulmoides* we identified only a partial E-domain sequence within the annotated sequence, while no G-domain sequence was identified.

**Data file S5:** MoeA Phylogenetic tree, obtained from maximum likelihood analysis. Bootstrap values are shown (pdf format).

**Data file S6:** Alignment file used for MoeA tree building (FASTA format)

**Data file S7:** Phylogenetic tree for MOCS2B obtained from maximum likelihood analysis (pdf format).

**Data file S8:** Phylogenetic tree for ADH obtained from maximum likelihood analysis (pdf format).

**Data file S9:** Local BLAST hits of MoeA homologs used for Alignment (FASTA format).
