## Supplementary figures and images for "Phylogenetic based dissection of eukaryotic Mo-insertase functionality: From mechanism to complex assembly"

### Data file S5

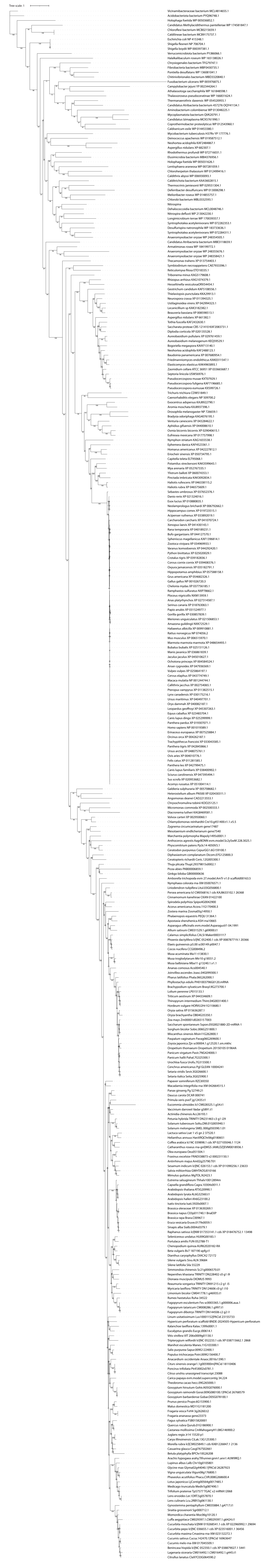

### Data file S7

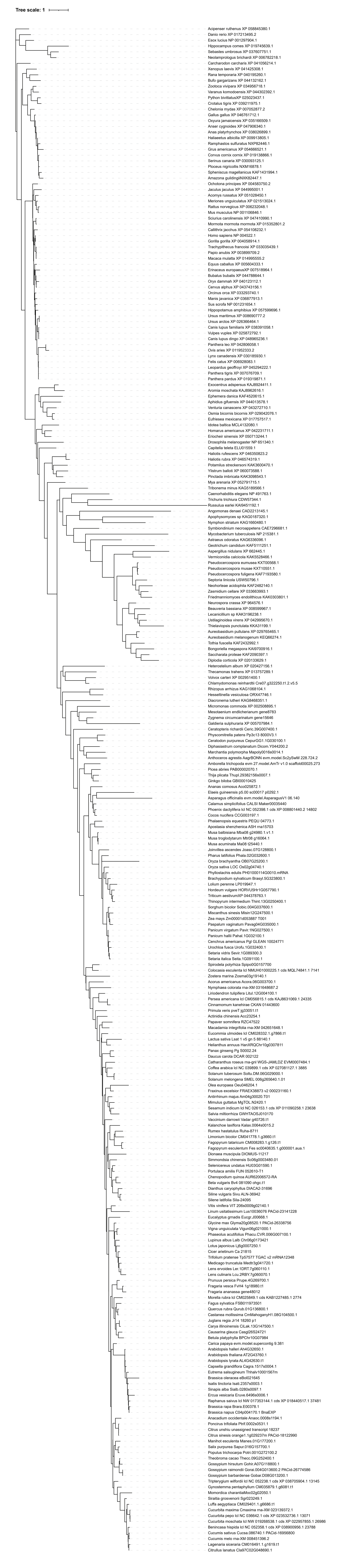

### Data file S7

#### ADH

Tree scale: 1

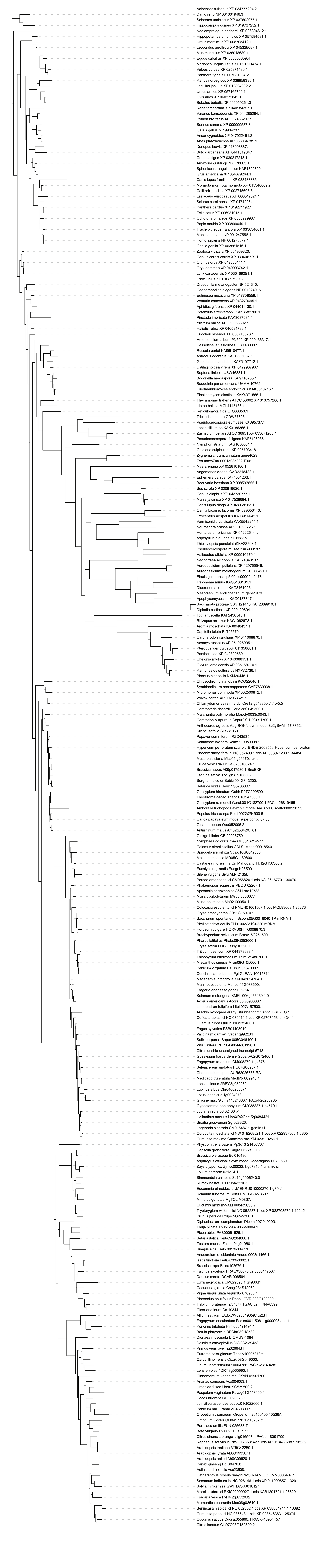
